# CAREPath: Semantic Context-Aware Reasoning Paths with Mechanism-Augmented Embeddings for Drug Repurposing

**DOI:** 10.64898/2026.06.09.731247

**Authors:** Haerin Song, Dongmin Bang, Bonil Koo, Sun Kim, Sangseon Lee

## Abstract

Biomedical knowledge graphs (BKGs) that include drugs, genes, and diseases support drug repurposing by connecting drugs to diseases through gene-mediated multi-hop paths, thereby enabling mechanism-of-action reasoning. However, deeper traversal does not necessarily improve mechanistic reasoning: long paths grow combinatorially and frequently pass through hub genes, producing irrelevant gene regulatory signals, whereas overly constrained or sparse paths may miss broader biological context. We propose CAREPath, a KG–LLM framework inspired by depth-first search (DFS)-like and breadth-first search (BFS)-like reasoning to balance mechanistic specificity, scalability, and context recovery. The DFS-like module constrains traversal to short disease–gene–drug paths, converts each path into a structured prompt, and encodes it with a biomedical language model to generate semantic path embeddings. Complementarily, the BFS-like module constructs entity-level mechanism-context embeddings from one-hop gene neighborhoods and enriches them through similarity-guided augmentation using pharmacologically related drugs and gene-signature-similar diseases. Across five biomedical KGs, CAREPath achieves the best overall AUPRC among 18 baselines, improving performance by up to 3.8%. Additional analyses show that semantic short-path encoding contributes most to performance, while mechanism-context augmentation improves robustness under sparse evidence and strengthens Gene Ontology functional agreement. Case studies and recently FDA-approved indications further demonstrate its practical relevance, positioning CAREPath as an interpretable framework for scalable and mechanism-aware drug repurposing. Source code is available at https://github.com/hamppy-song/CAREPath.

## Introduction

Drug repurposing, the identification of new indications for existing drugs, remains a central challenge in systems pharmacology [1]. Success requires not only ranking candidate drug–disease pairs but also explaining the mechanisms that connect them. Many diseases involve molecular disruptions (e.g., dysregulated gene expression) that drugs can modulate through their mechanisms of action [2]. Biomedical knowledge graphs (BKGs) help model these mechanisms by integrating heterogeneous entities (e.g., genes, diseases, and drugs) and their relationships [3]. A common strategy is path-based reasoning, which links drugs and diseases through multi-hop paths over intermediate gene nodes, using heterogeneous walks or by linearizing KG facts into sequences for representation learning [5]. Such paths can represent routes by which drug-target interactions propagate through gene networks to affect disease processes [6]. Since disease-relevant regulation is often distributed across several genes rather than a single target, such gene-mediated paths offer a natural substrate for mechanism-of-action reasoning [7].

However, increasing hop depth does not necessarily improve mechanistic reasoning in KG-based drug repurposing. Many existing approaches rely on multi-hop traversal to capture indirect biological relationships, assuming that deeper exploration provides richer evidence. In practice, however, this assumption introduces two challenges.

First, deeper traversal substantially increases the number of candidate paths and frequently routes reasoning through hub genes that connect many unrelated biomedical contexts [8, 9]. As path complexity grows, representations derived from counts [10], walks [11], or proximity-based [12] features can increasingly reflect graph connectivity rather than biologically meaningful mechanisms. Although such paths remain topologically valid, they often become pair-nonspecific and dilute the compact gene-mediated signals that are most informative for drug– disease reasoning [13]; we further examine this phenomenon through hub-gene and path-specificity analyses (Supplementary Section S5).

Second, reducing traversal complexity alone is insufficient because constrained or sparse paths may fail to capture broader mechanism context required for robust repurposing [14]. Short disease–gene–drug paths provide interpretable and pair-specific evidence, but they may be absent, incomplete, or dominated by a small number of generic mediators when the KG lacks sufficient annotations for a drug or disease [15]. This is especially limiting for sparsely annotated entities, motivating the use of related drugs and diseases as a complementary source of mechanism context. Thus, effective KG-based repurposing requires not only controlling noisy multi-hop expansion but also recovering missing mechanism-level context in a biologically interpretable way.

To address these challenges, we propose CAREPath (Context-Aware REasoning Path), a KG–LLM framework that balances mechanistic specificity, scalability, and context recovery in drug repurposing. CAREPath first applies a DFS-inspired path reasoning strategy that constrains traversal to short disease– gene–drug paths, limiting the number of intermediate genes to suppress combinatorial path expansion and reduce hub-gene-driven paths that recur across many unrelated drug–disease pairs. Rather than treating these paths only as topological patterns, CAREPath converts each constrained path into a structured natural-language prompt and encodes it with a biomedical language model, thereby preserving mechanistic semantics within compact gene-mediated traces. Because short paths may still be sparse or incomplete, CAREPath further incorporates a BFS-inspired mechanism-context module that constructs entity-level embeddings from one-hop gene neighborhoods and enriches them through similarity-guided augmentation using pharmacologically related drugs and gene-signature-similar diseases. The resulting representation combines pair-specific semantic path information with broader mechanism-level context for drug–disease association prediction.

Across five biomedical KG benchmarks (MSI [16], PrimeKG [17], Hetionet [10], SuppKG [18], and KEGG50k [19]), CAREPath achieves the best overall AUPRC against 18 baselines, even in the disease cold-start setting, with gains of up to 3.8%. Additional analyses further show that the gains arise from semantically encoded short-path reasoning and mechanism-context augmentation, which together improve robustness under sparse evidence settings and strengthen biological interpretability. Case studies and evaluation on recently FDA-approved indications further support the practical relevance of the framework. Together, these findings position CAREPath as an interpretable KG–LLM framework that reduces noisy long-range traversal while recovering missing mechanism-level context.

## Materials and Methods

In this section, we introduce CAREPath, a drug–disease association prediction framework that encodes drug–disease regulatory mechanisms using two complementary embeddings (Fig. 1). In the DFS module, CAREPath extracts constrained disease–gene–drug paths and encodes templated natural-language prompts with a biomedical language model to obtain semantic path embeddings. The BFS module constructs mechanism-context embeddings for the drug and disease from one-hop gene neighborhoods via similarity-guided pooling (drug class-based for drugs and gene-signature-based for diseases). CAREPath then aggregates path embeddings via max pooling and fuses them with the mechanism-context embeddings, which are fed into a classifier for the final drug–disease association prediction.

**Fig. 1.**
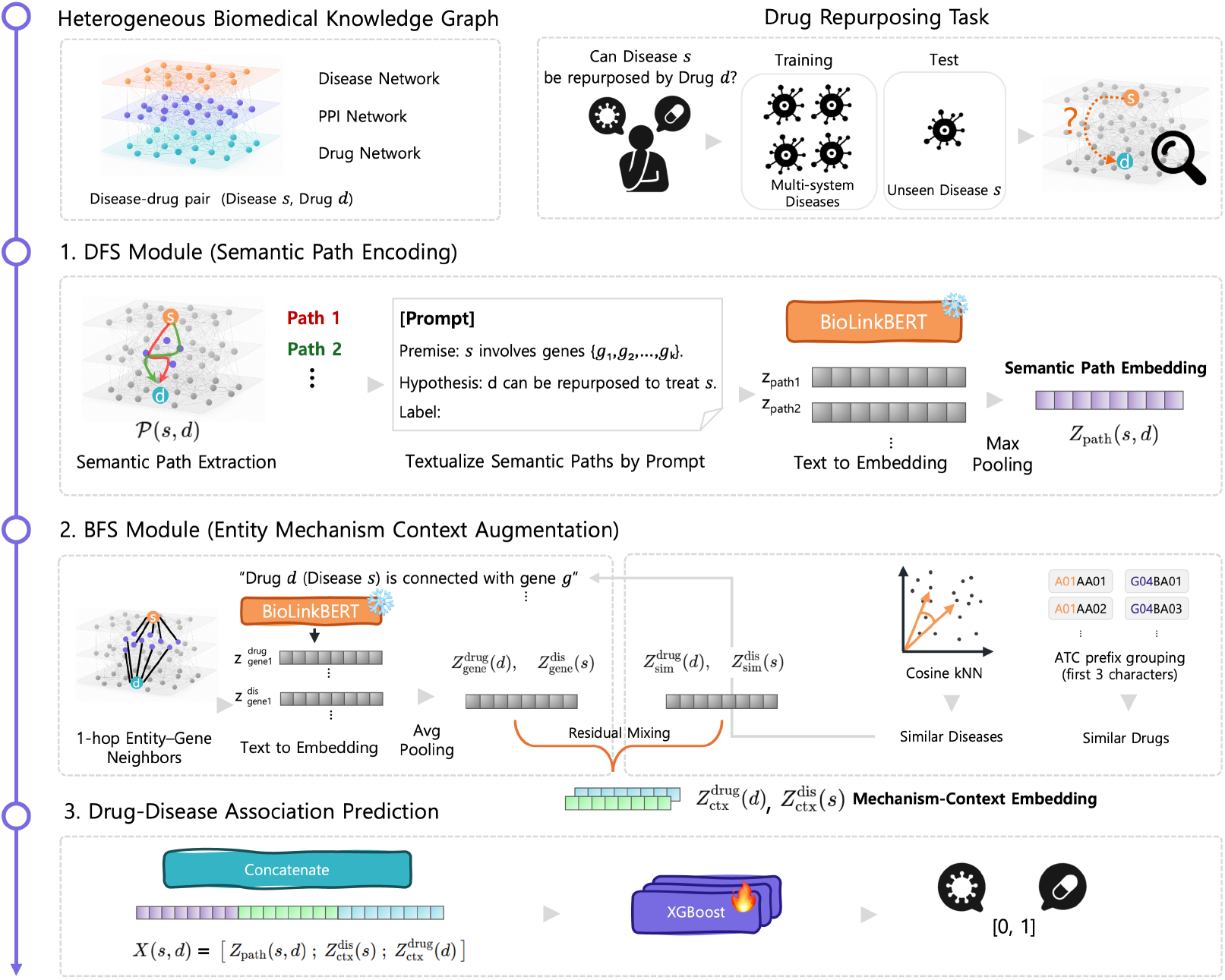
Overview of the CAREPath framework. Given a disease–drug pair (*s, d*), CAREPath extracts constrained disease–gene–drug paths from a heterogeneous knowledge graph, converts each path into a structured prompt, and encodes it with BioLinkBERT; max pooling aggregates these into a pair-specific path embedding. In parallel, CAREPath builds mechanism-context embeddings from the 1-hop gene neighborhoods of the drug and disease and enriches them via similarity-guided pooling (cosine kNN for diseases; ATC-prefix-related drugs) with residual mixing. The final representation concatenates the path embedding and the two mechanism-context embeddings, and is scored by a stacking ensemble of XGBoost classifiers to predict the drug–disease association.

### Problem Formulation

We consider drug–disease association prediction on a BKG 𝒢 = (𝒱, *ε*) with node set 𝒱 = 𝒱_drug_ ∪ *𝒱*_disease_ ∪ *𝒱*_gene_, where 𝒱_gene_ may include genes and proteins depending on the KG. Edges *E* represent typed biomedical relations (e.g., gene–gene, drug–gene, and disease–gene). Drug–disease associations are given as labeled pairs 𝒟 = *{*(*s, d, y*_*s,d*_) | *s ∈ 𝒱*_disease_, *d ∈ 𝒱*_drug_, *y*_*s,d*_ *∈ {*0, 1*}}*. Note that these associations are not explicitly represented as edges in 𝒢.

Given a pair (*s, d*), the goal is to learn a predictor *f* that estimates the likelihood of an association, *ŷ*_*s,d*_ = *f* (*s, d* | *G*) *∈* [0, 1], trained such that *ŷ*_*s,d*_ matches the true label *y*_*s,d*_.

### Constrained Semantic Path Encoding (DFS module) *Path Extraction from BKG*

To represent candidate regulatory mechanisms linking a disease and a drug, we extract constrained disease–gene–drug paths from the BKG 𝒢 = (𝒱, *ε*). For each disease *s ∈ 𝒱*_disease_ and drug *d ∈ 𝒱*_drug_, we enumerate simple paths from *s* to *d* subject to a constraint on the number of intermediate gene nodes, reflecting short mechanistic chains in which disease-associated genes mediate therapeutic targets. Concretely, we retain only paths containing at most *k*_max_ intermediate gene nodes. We set *k*_max_ = 2 based on a coverage–redundancy trade-off analysis across BKGs (Details in Supplementary Section S5). These constraints control the growth of candidate paths in dense BKGs and reduce redundancy driven by highly connected nodes, while prioritizing compact, mechanism-oriented traces.

Under these constraints, some drug–disease pairs may yield no valid paths due to sparsity or missing links in the graph. To increase coverage without relaxing the hop limit, inspired by DREAMwalk [4], we apply a drug class-guided drug substitution strategy: when no path is found between (*s, d*), we select a pharmacologically related substitute drug *d*^*′*^ from the same Anatomical Therapeutic Chemical (ATC) code group as *d* (restricted to drugs present in 𝒢) and re-run constrained path extraction for (*s, d*^*′*^). We then use the resulting gene set as a proxy mechanism to construct path prompts for the original pair (*s, d*).

### Prompt Template Construction

Given the extracted path set *P*_*s,d*_, we convert each path into a structured natural-language prompt for semantic encoding with a biomedical language model. For each path *p*_*i*_ *∈ P*_*s,d*_, we construct a prompt *x*_*i*_ using a natural language inference (NLI)-style template:

**Premise:** *s* involves genes *{g*_1_, …, *g*_*k*_*}*.

**Hypothesis:** *d* can be repurposed to treat *s*.

**Label:**

This template yields compact, mechanism-oriented inputs that summarize the genes implicated by each constrained path while keeping the drug–disease query explicit in the hypothesis. When no valid path is available even after ATC-based substitution, we use the same template with an empty gene set (i.e., genes none), ensuring all drug–disease pairs produce well-formed prompts under a uniform input format.

### Semantic Path Embedding

To encode the prompts, we use a domain-specific biomedical language model, BioLinkBERT [20], as a frozen sentence encoder *ϕ*(*·*). For each prompt *x*_*i*_, we obtain a dense representation by extracting the final-layer [CLS] embedding: **z**_*i*_ = *ϕ*(*x*_*i*_) *∈ ℝ*^*D*^, where *D* is the hidden dimension. For a given drug–disease pair (*s, d*), this yields a set of semantic path embeddings 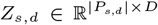
, whose rows correspond to *{***z**_1_, …, **z**_|*P*_*s,d*_|_*}*. Heterogeneous To summarize multiple candidate mechanisms into a single pair-level representation, we apply element-wise max pooling: **Z**_path_(*s, d*) = MaxPool(*Z*_*s,d*_) *∈ ℝ*

### Entity Mechanism-Context Augmentation (BFS module)

#### Gene Neighborhood Retrieval

In addition to path-level representations, CAREPath constructs *mechanism-context embeddings* for each disease and drug from their local gene neighborhoods in the BKG. Short mechanistic paths provide specific, interpretable signals, but they are often missing or unreliable when KG neighborhoods are sparse or hub-dominated. To provide a robust entity-level prior, CAREPath therefore complements DFS path traversal with a BFS expansion that summarizes each entity’s local gene neighborhood. This context is designed to remain informative even when pair-specific path signals are limited. For any node *x ∈ 𝒱*_drug_ ∪*𝒱*_disease_, we retrieve its one-hop gene neighborhood

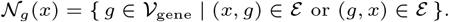

These neighborhoods provide a compact, entity-centric summary of genes associated with the disease (e.g., dysregulated genes) and genes linked to the drug (e.g., targets or affected genes). We retain all gene neighbors in 𝒱_*g*_(*x*) without subsampling, as incorporating the BFS-derived mechanism-context improves performance across all density strata (detailed analysis provided in Supplementary Section S6). Mean pooling over sentence embeddings yields a fixed-dimensional context vector that is invariant to neighborhood size, so highly connected entities do not require explicit truncation.

#### Neighborhood Sentence Construction

We convert each entity’s one-hop gene neighborhood into a set of natural-language sentences for semantic encoding with a biomedical language model. For each drug *d* (or disease *s*), we retain only gene-type one-hop neighbors to avoid leakage from drug–disease co-neighborhoods. We use these sentences as lightweight textual surrogates of local functional programs, enabling the biomedical language model to encode neighborhood-level semantics in a shared embedding space. Each retained neighbor *g* is mapped to a human-readable name and converted into a sentence describing local mechanistic context:

*t*(*d, g*) = Drug *d* is connected to *g. t*(*s, g*) = Disease *s* is connected to *g*.

*N*_*g*_(*x*) = *{ g ∈ V*_gene_ | (*x, g*) *∈ E* or (*g, x*) *∈ E}*.

Let *T* (*d*) = *{t*(*d, g*) | *g ∈ 𝒱*_*g*_(*d*)*}* and *T* (*s*) = *{t*(*s, g*) | *g ∈ 𝒱*_*g*_(*s*)*}* denote the resulting sets of neighborhood sentences. If 𝒱_*g*_(*x*) is empty, we use a single fallback sentence (i.e., Drug *d* is `connected to genes none`.) so that |*T* (*v*)| = 1 and the encoding step remains well-defined.

#### Mechanism-Context Embeddings and Similar-neighbor Augmentation

We encode each neighborhood sentence using the biomedical language model encoder *ϕ*(*·*) and compute an initial mechanism-context embedding by mean pooling over the sentence set. For a drug *d* and disease *s*,

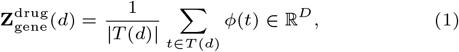

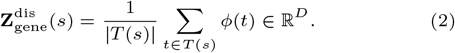

To improve robustness when one-hop neighborhoods are sparse or noisy, we apply similar-neighbor augmentation via residual mixing. Let 𝒦^drug^(*d*) denote a set of pharmacologically related drugs (ATC-based) and 𝒦^disease^(*s*) denote a set of similar diseases (gene-signature-based). We first summarize neighbors with a pooling operator Pool(*·*) (mean or similarity-weighted mean), and then form augmented embeddings:

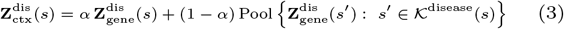

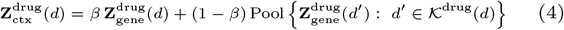

with *α, β ∈* [0, 1]. We use these augmented mechanism-context embeddings together with semantic path embeddings to balance specific mechanistic information and broader functional compatibility. Details of the similarity definitions and pooling are provided in Supplementary Section S2.

### Feature Fusion and Association Prediction

For each drug–disease pair (*s, d*), CAREPath constructs a unified feature vector by concatenating (i) the pair-specific semantic path embedding from constrained mechanistic paths (Section 2.2) and (ii) the entity-level mechanism-context embeddings for the drug and disease (Section 2.3):

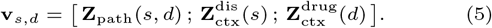

We use **v**_*s,d*_ as input to a supervised classifier to estimate the association probability *ŷ*_*s,d*_ = *f* (**v**_*s,d*_) *∈* [0, 1].

We instantiate *f* (*·*) with a stacking ensemble, which is well-suited for heterogeneous concatenated feature vectors and provides competitive performance with limited additional modeling assumptions. Concretely, three XGBoost models with random seeds *{r*_0_, *r*_0_ + 1, *r*_0_ + 2*}* (where *r*_0_ is the base seed) serve as base learners, each trained independently with a logistic objective for binary classification. Their out-of-fold predicted probabilities are then combined by a logistic regression meta-learner to obtain the final score *ŷ*_*s,d*_.

### Datasets and Evaluation Setups

(1) We evaluated CAREPath on five BKG benchmarks: MSI [16], PrimeKG [17], Hetionet [10], SuppKG [18], and KEGG50k [19]. To ensure comparability, each BKG was preprocessed to retain

(2) the node and relation types relevant to gene-mediated drug– disease reasoning. Dataset details and preprocessing are provided in Supplementary Section S3.

To assess the generalization ability of the model across diverse data distributions, we employed 5-fold cross-validation under three split settings: random, disease cold-start, and drug cold-start. Random splits were label-stratified, while cold-start splits were group-based using GroupKFold, where the grouping key is the individual disease (for disease cold-start) or drug (for drug cold-start) entity. This guarantees that all pairs involving a given disease or drug appear in a single fold, so that test entities are entirely unseen during training.

The base-learner hyperparameters were tuned only once, in a separate stage that was kept independent of the reported evaluation. This tuning used disease-grouped splitting: we performed a randomized hyperparameter search on a held-out group split and re-evaluated the best candidates with disease-grouped 5-fold cross-validation, selecting the most stable configuration. These hyperparameters were then fixed and used unchanged across all three split settings (random, disease cold-start, and drug cold-start) and all evaluation folds. As a result, no hyperparameter tuning was performed on the evaluation data, avoiding information leakage into the reported results. The meta-learner was trained using inner 5-fold cross-validation within each training fold. Models were evaluated using accuracy, AUROC, AUPRC, and F1. Experimental details are provided in Supplementary Section S4.

## Results

### Performance Comparison on Five BKGs

We benchmark CAREPath against a diverse set of baselines grouped into four families: **Graph-based** GNNs (GraphSAGE [21], CompGCN [22], GIN [23], GAT [24]), **KGE-based** models (TransE [25], TransR [26], RotatE [27], ComplEx [28], RESCAL [29]), **LLM-based** text encoders (BioBERT [30], BioLinkBERT [20], PubMedBERT [31], SapBERT [32]), and **Path-based** reasoning methods (Node2Vec [11], Drugrep-KG

[5], DREAMwalk [4], FuseLinker [33], K-Paths [34]). Details of baseline models are described in Supplementary Section S1. Experiments were performed on five BKGs (MSI, PrimeKG, Hetionet, SuppKG and KEGG50k) under three evaluation settings: *random split, disease cold-start* and *drug cold-start*. We use AUPRC as the primary metric because it directly reflects retrieval quality when prioritizing candidate drug– disease associations. Full results for all metrics across the three settings are provided in the Supplementary Section S7 (Tables S4–S18).

Under the disease cold-start split, CAREPath achieves the highest AUPRC on all five KGs (Table 1), including MSI (0.893) and Hetionet (0.965). By contrast, baseline performance varies substantially across KGs, with no method consistently ranking near the top. Several recent baselines, including DREAMwalk, DrugRep-KG, FuseLinker, and K-Paths, achieve strong AUPRC on one or two KGs but drop substantially on others. For instance, K-Paths reaches 0.907 on Hetionet but drops to 0.503 on KEGG50k, while FuseLinker reaches 0.927 on Hetionet but only 0.806 on MSI. Notably, CAREPath also outperforms standalone BioLinkBERT—the same biomedical language model used as its encoder—across all five KGs (e.g., MSI: 0.893 vs 0.849; PrimeKG: 0.944 vs 0.878; Hetionet: 0.965 vs 0.924). This indicates that the gains stem from the structural design of CAREPath rather than from the language model alone.

**Table 1.**
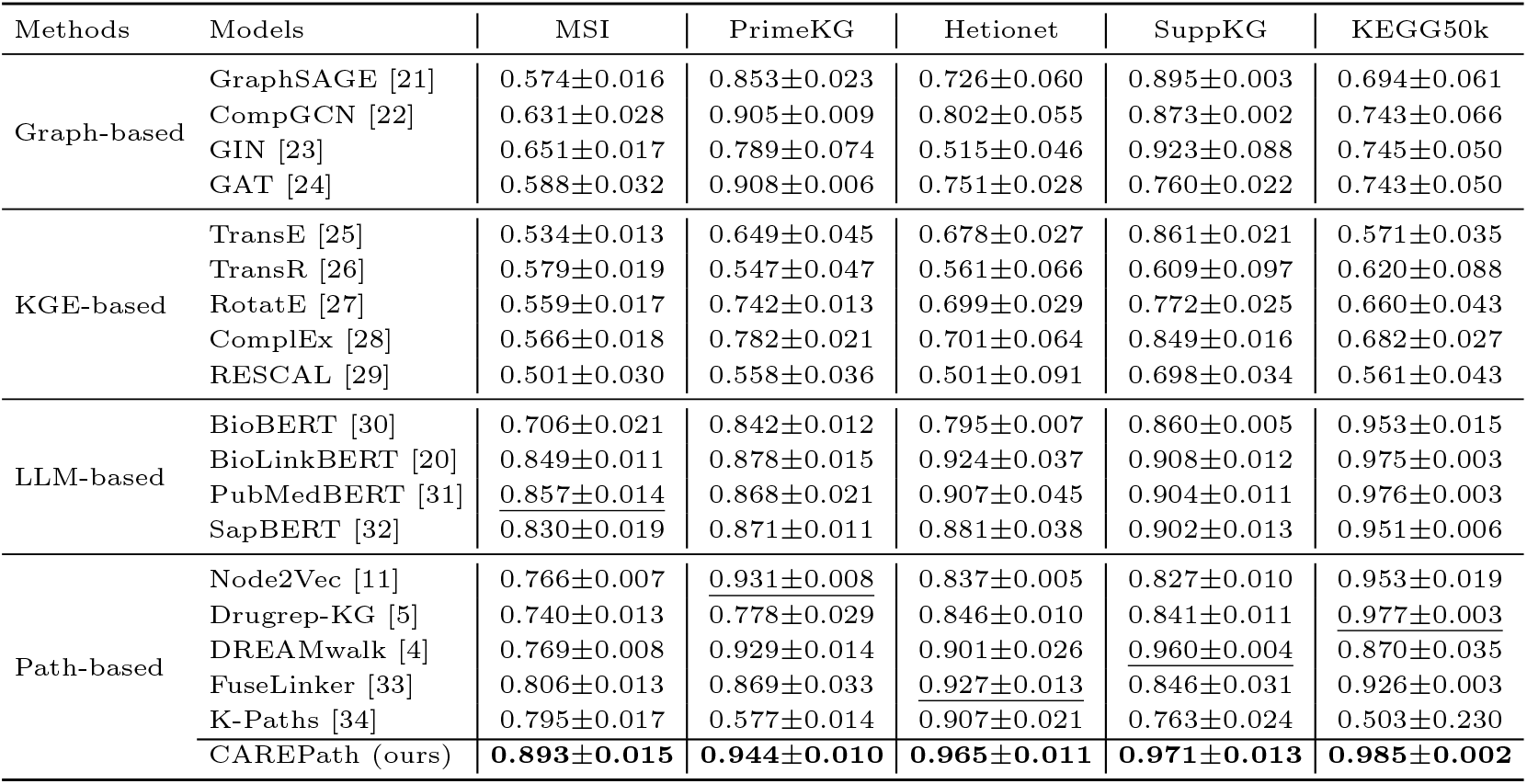
Performance comparison (AUPRC) across five biomedical knowledge graphs under the disease cold-start split. Results are reported as mean *±* standard deviation over five folds. Baselines are grouped by modeling paradigm.

Taken together, these results suggest that proximity-driven models and unconstrained multi-hop reasoning are sensitive to KG-specific relation patterns and connectivity. CAREPath mitigates this sensitivity by combining constrained short-path reasoning with semantic path encoding and BFS-like mechanism-context augmentation, thereby preserving pair-specific gene-mediated signals while compensating for sparse or incomplete path evidence. Similar trends are observed under the random split and the drug cold-start split (Supplementary Section S7).

### Effect of Semantic Path Embedding and Mechanism-Context Representations

We evaluated the contribution of the two modules in CAREPath:

(1) semantic path embedding and (2) mechanism-context representations, on MSI under the disease cold-start split (Fig. 2). The full model achieves a mean AUPRC of 0.893, and ablating semantic path embedding causes the largest decrease (*−*4.6%; paired t-test, *p* = 2.994 *×* 10^*−*4^), indicating that path-level semantics provide the dominant signal for gene-mediated drug–disease reasoning. Excluding the drug or disease mechanism-context embedding yields smaller but consistent reductions *−*1.1% and *−*1.5%, respectively, and removing both further decreases performance by *−*2.3%. These trends suggest that mechanism-context representations supply complementary mechanistic information, whereas semantic path encoding plays a key role in preserving mechanistic meaning under constrained short-path reasoning.

**Fig. 2.**
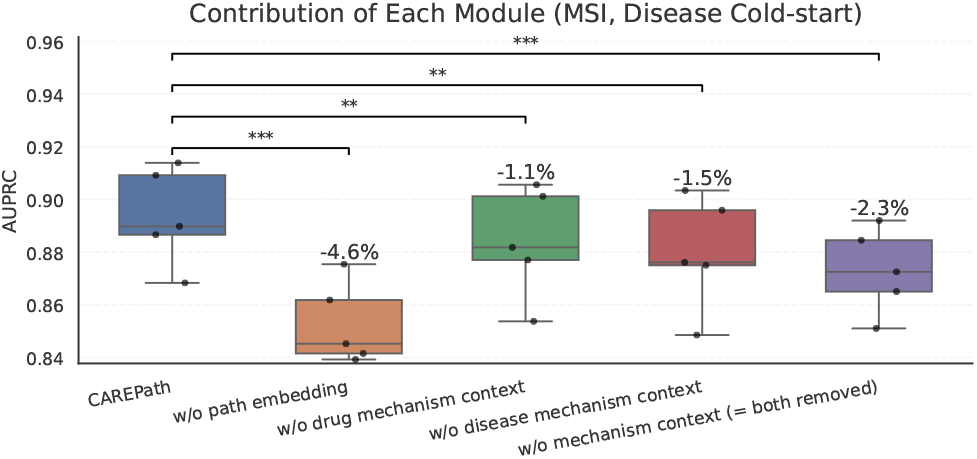
Ablation analysis of CAREPath modules on MSI under the disease cold-start setting. Ablating the semantic path embedding yields the largest performance drop, while removing the drug or disease mechanism-context embedding results in smaller but consistent reductions. (*: *p <* 0.05, **: *p <* 0.01, ***: *p <* 0.001, paired t-test).

### Disease-level Mechanistic Similarity in Semantic Path Embeddings under Short-Path Constraint

We further examined whether semantic path embeddings learned from constrained short paths capture meaningful mechanistic structure using a hypothesis-driven case set. Because such short paths suppress hub-driven generic routes and emphasize gene mediators specific to each disease–drug pair, we expect the resulting embeddings to organize diseases according to their underlying regulatory programs rather than topological proximity alone. We selected two mechanistically related lymphoid malignancies: Hodgkin disease and acute lymphocytic leukemia. We included thrombosis as a mechanistically distinct comparator to test whether the embedding space groups related diseases while separating an unrelated condition. We computed disease-level embeddings on MSI from short disease– gene–drug paths encoded by a biomedical language model and visualized them with principal component analysis (PCA) (Fig. 3). To isolate the effect of short-path semantic encoding, we used semantic path embeddings only, without entity-context augmentation.

**Fig. 3.**
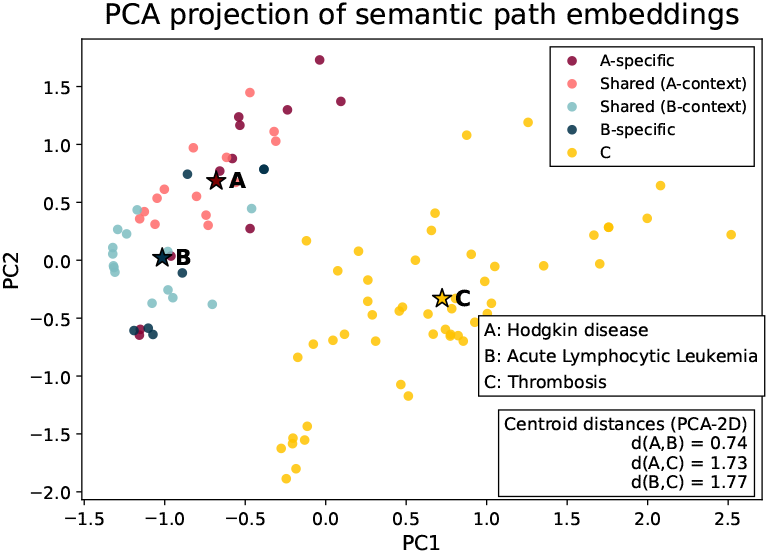
PCA of semantic path embeddings. Related diseases (A: Hodgkin disease, B: Acute lymphocytic leukemia) cluster together, while a distinct disease (C: thrombosis) is separated.

As shown in Fig. 3, Hodgkin disease (A) and acute lymphocytic leukemia (B) cluster closely, whereas thrombosis (C) is well separated. Within the A-B region, the overlap reflects common lymphoid programs [35], while disease-specific points capture microenvironment-dominated regulation in Hodgkin disease versus leukemia-intrinsic proliferation in Acute lymphocytic leukemia. [36]. Thrombosis is primarily driven by coagulation and vascular inflammation pathways [37], consistent with its separation from the lymphoid cluster. These results suggest that semantic encoding preserves mechanistic signals, clustering related diseases while separating distinct conditions even under short-path constraints.

### Robust Performance under Abundant and Sparse Entity Neighborhoods

In BKGs, drugs and diseases vary substantially in how richly they are annotated with gene associations. Some entities have rich 1-hop gene neighborhoods that provide ample mechanistic context, whereas others are sparsely annotated, limiting the available signals for gene-mediated drug–disease reasoning. To test whether CAREPath remains reliable across these varying connectivity levels on MSI, we stratified drug–disease pairs by the size of the 1-hop gene neighborhoods of the drug and the disease and compared performance against recent baselines.

For each pair, we counted genes directly linked to the drug and disease nodes, and defined two subsets: (i) *abundant* pairs when at least one entity falls in the top 15% by linked-gene count (more than 80 linked genes) and (ii) *sparse* pairs when at least one entity has fewer than two linked genes. This analysis includes 1,822 abundant pairs and 4,412 sparse pairs.

Across CAREPath and recent baselines (DREAMwalk, FuseLinker, and K-Paths), performance is higher on abundant pairs than on sparse pairs, consistent with the greater availability of gene-level mechanistic reasoning in well-annotated neighborhoods (Fig. 4a). CAREPath achieves the highest AUPRC in both strata and retains a clear advantage under sparse neighborhoods, where all methods experience performance degradation (Fig. 4b). This pattern suggests that CAREPath is less dependent on dense local annotations than the competing path-based methods and remains effective when direct gene-mediated regulation is limited.

**Fig. 4.**
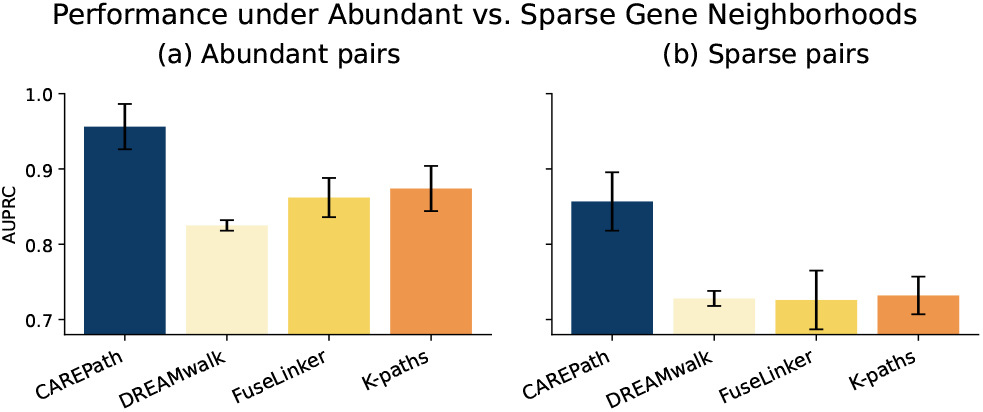
Robustness under abundant and sparse entity neighborhoods: AUPRC on (a) abundant pairs and (b) sparse pairs across CAREPath and recent baselines. CAREPath remains competitive when gene neighborhoods are abundant and exhibits the smallest degradation under sparsity, indicating reduced sensitivity to sparsely-annotated entities.

### Improved GO Functional Agreement with Mechanism-Context Embeddings

We evaluated on MSI whether the mechanism-context embeddings improve functional agreement with reference Gene Ontology biological process (GO-BP) annotations beyond what is captured by the semantic path embedding alone. Because constrained short paths can be missing or incomplete for many drug–disease pairs, we compared two variants of CAREPath: a DFS-only variant and a DFS+BFS variant. For each variant, we identified enriched GO-BP terms from the implicated gene set using Enrichr [38], and computed an inverse document frequency (IDF)-weighted Jaccard similarity against reference GO-BP annotations derived from external resources (DrugBank [39] for drugs; DisGeNET [40] for diseases), with details provided in Supplementary Section S8.

As shown in Fig. 5, adding the mechanism-context embeddings improves both agreement and recall over reference GO-BP annotations, with a median recall gain of +24.5% relative to the DFS-only variant. The gains are largest when only a few DFS-derived GO terms are available and gradually diminish as the number of available terms increases. This pattern suggests that mechanism-context embeddings provide complementary functional information when short-path signals are sparse, while adding less new information once DFS-derived functional coverage is already sufficient.

**Fig. 5.**
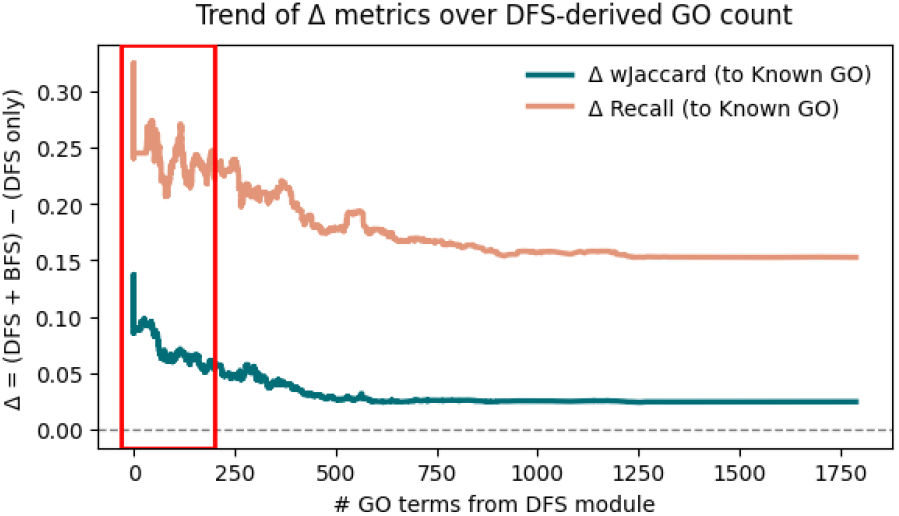
Improvement in agreement (Δ IDF-weighted Jaccard) and recall (Δ Recall) over reference GO-BP annotations versus the number of DFS-derived GO terms (DFS+BFS vs. DFS-only). The red box highlights the sparse regime where mechanism-context embeddings contribute the largest gains.

### Explainable Drug Repurposing: Recovering Mechanistic Context Beyond Constrained Paths

To examine how the BFS-derived mechanism-context complements constrained path-based reasoning, we compared the intermediate genes and associated KEGG pathways retrieved by the constrained path module alone (DFS-only) against those retrieved by the full CAREPath framework. The full framework augments the DFS-only paths with the BFS-derived mechanism-context, which summarizes each target’s own one-hop connected genes and then broadens this neighborhood by pooling pharmacologically similar drugs (grouped by ATC code) and gene-signature-similar diseases. We present two drug–disease pairs from therapeutic areas in which efficacy is typically described as involving multiple receptor families or cross-pathway signaling: oncology (Capecitabine *→* Mammary Neoplasms) and psychiatry (Doxepin *→* Bipolar Depression).

For each pair, we report the KEGG pathways retrieved by each module together with reported associations from the published literature.

### Case 1: Capecitabine → Mammary Neoplasms

For Capecitabine, the constrained path (DFS-only) module connected the drug to the disease through only a single gene mediator, *TYMS*, within the Pyrimidine metabolism pathway (Fig. 6a). This correctly reflects Capecitabine’s primary mechanism as a thymidylate-synthase-inhibitor prodrug that blocks DNA synthesis [41]; however, this represents a generic antimetabolite mechanism shared across many cancers and does not include ER, PR, and HER2, clinically used biomarkers for subtype classification and therapeutic stratification in breast cancer [42].

**Fig. 6.**
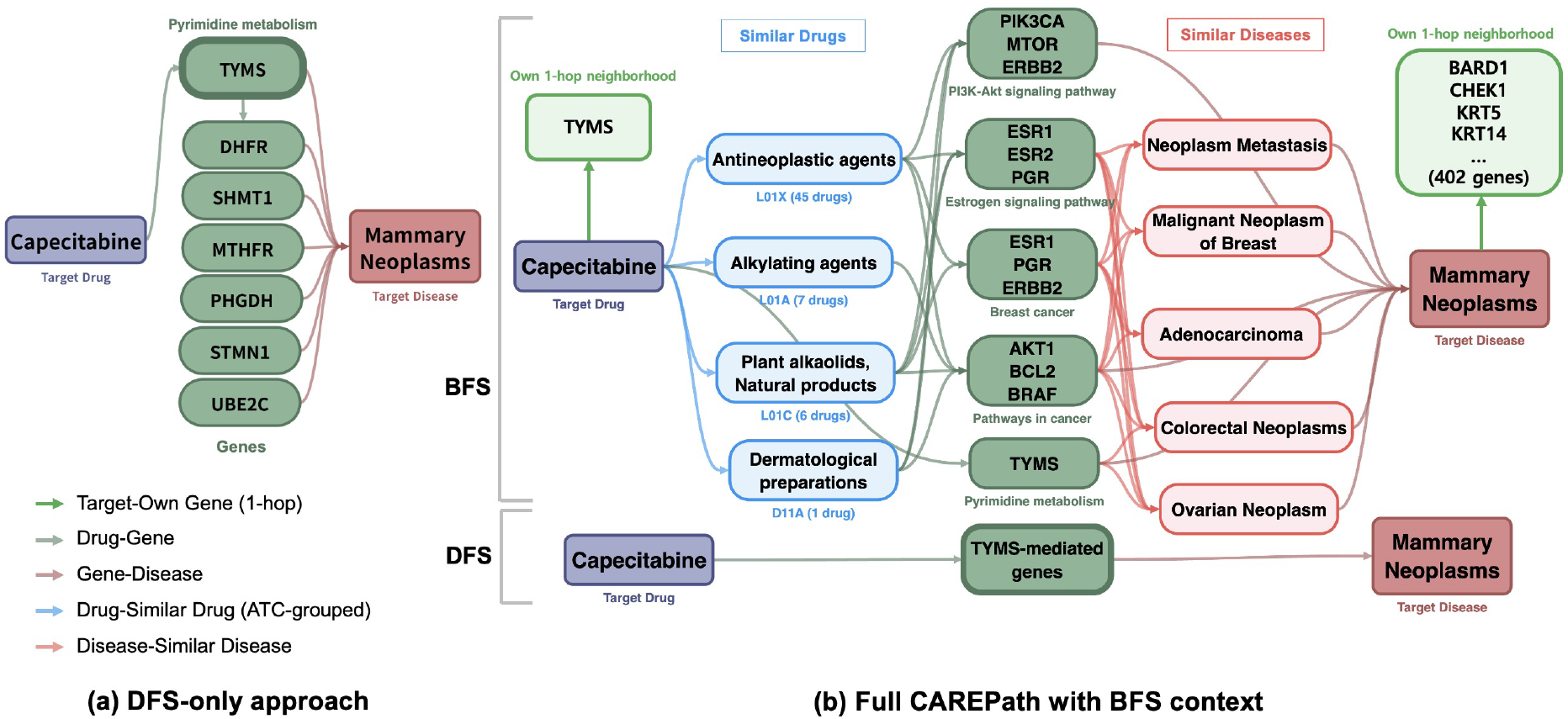
Recovering breast-cancer-specific mechanistic context for Capecitabine *→* Mammary Neoplasms. **(a)** The DFS-only module connects the drug to a single target gene (*TYMS*, Pyrimidine metabolism), capturing only the generic DNA-synthesis-blockade mechanism that is not specific to mammary neoplasms. **(b)** The full CAREPath first incorporates each target’s own one-hop gene neighborhood (BFS core): the disease (Mammary Neoplasms) contributes breast-cancer-relevant genes such as *BARD1, CHEK1, KRT5*, and *KRT14* that are not recovered through similar-neighbor augmentation, whereas the drug (Capecitabine) connects to only a single gene (*TYMS*). Because the drug’s own neighborhood is too sparse on its own, CAREPath augments it by traversing ATC-grouped similar drugs and gene-signature-similar diseases, additionally retrieving breast-cancer-relevant pathways including *PI3K-Akt signaling* (antimetabolite resistance) and *Estrogen signaling* (hormone-dependent growth). The BFS core and similar-neighbor augmentation thus play complementary roles. Intermediate genes are grouped by KEGG pathway membership.

The BFS module first incorporates each target’s own one-hop gene neighborhood. Here the two targets are markedly asymmetric: Mammary Neoplasms is richly annotated (402 connected genes, including breast-cancer-relevant genes such as *BARD1, CHEK1, KRT5*, and *KRT14* that are not recovered through similar-neighbor augmentation), whereas Capecitabine connects to only a single gene (*TYMS*). Because the drug’s own neighborhood is too sparse to recover breast-cancer-specific context on its own, the full CAREPath framework further traverses similar drugs (ATC-grouped antineoplastic agents) and gene-signature-similar diseases (e.g., Neoplasm Metastasis, Malignant Neoplasm of Breast), additionally retrieving the PI3K– Akt signaling pathway, containing survival mediators including *PIK3CA, MTOR*, and *ERBB2*. PI3K–Akt activation is a well-characterized bypass route through which tumor cells maintain proliferation under antimetabolite pressure, and combination therapy with PI3K inhibitors such as Alpelisib has been investigated as a strategy to address PI3K-pathway-mediated chemoresistance [43]. CAREPath also retrieved the Estrogen signaling pathway, reflecting the bidirectional crosstalk between PI3K and ER signaling that sustains hormone-dependent tumor growth and underlies endocrine-therapy resistance [44]. This case illustrates the complementary roles of the two BFS components: the disease’s own neighborhood contributes specific genes invisible to augmentation, while augmentation compensates for the drug’s sparse neighborhood. Together, these neighborhoods place the cytotoxic action of Capecitabine within the regulatory context that governs treatment response in breast cancer subtypes.

### Case 2: Doxepin → Bipolar Depression

For Doxepin, the constrained path (DFS-only) module retrieved two serotonergic gene mediators (*HTR1A, SLC6A4*) within the Serotonergic synapse pathway (Supplementary Figure S3). This reflects the classical monoamine hypothesis on which tricyclic antidepressants such as Doxepin were originally developed [45]; however, this view alone does not capture the synaptic-balance and neuroplasticity mechanisms that contemporary research has implicated in the pathophysiology of bipolar depression and in its treatment response [46].

The BFS module first incorporates each target’s own one-hop gene neighborhood. Unlike the sparse drug neighborhood in Case 1, Doxepin is itself connected to a broad set of receptor genes spanning serotonergic (*HTR1A, HTR2A*), adrenergic (*ADRA* family), muscarinic (*CHRM* family), and histaminergic (*HRH* family) systems, consistent with its multi-receptor pharmacology, while Bipolar Depression itself contributes mood-disorder-related genes such as *COMT, DISC1, TPH2*, and *GRIK4*. Building on these neighborhoods, the full CAREPath framework further traverses similar drugs (ATC-grouped antidepressants, psychostimulants, psycholeptics, anti-dementia drugs, and antiobesity preparations) and gene-signature-similar diseases (e.g., Bipolar disorder, Depressive disorder), additionally retrieving the Neuroactive ligand-receptor interaction pathway, which contained glutamatergic (*GRIN* family), GABAergic (*GABRA*/*GABRB* family), and adrenergic (*ADRA* family) receptor genes. Dysregulation of excitatory/inhibitory balance has been reported as a key pathological feature of mood disorders, and restoring this balance is the rationale behind glutamatergic agents such as ketamine in treatment-resistant depression [47]. CAREPath also retrieved components of the cAMP signaling pathway, a downstream cascade mediating BDNF-dependent neuroplasticity that has been proposed as a convergent mechanism of antidepressant action [48]. Finally, CAREPath retrieved dopaminergic components beyond the serotonergic neighborhood; this is consistent with reports that mood-state transitions in bipolar depression involve integrated dopamine-serotonin receptor signaling rather than serotonergic modulation alone [49]. This case illustrates how the two BFS components progressively broaden the DFS-only view: the target’s own one-hop neighborhood already extends beyond serotonin to multiple receptor systems, while similar-neighbor augmentation further recovers downstream synaptic and neuroplasticity pathways not reachable from the short path alone. Together, these neighborhoods recover the broader synaptic and neuroplasticity mechanisms now implicated in contemporary depression treatment, situating Doxepin’s classical monoaminergic action within this wider neuro-modulatory context.

### Evaluation on Newly FDA-approved Indications

To demonstrate real-world repurposing relevance beyond KG-derived evaluation splits, we evaluated trained models on drug– disease pairs corresponding to new FDA-approved indications (label expansions) in 2024–2025^1^. This set provides an external assessment of whether model scores prioritize associations that were later approved in clinical practice. We restricted evaluation to drug and disease entities covered by our BKG and removed any pairs overlapping with training data, leaving eight previously unseen drug–disease associations (Supplementary Section S9).

Table 2 compares CAREPath with representative baselines. CAREPath assigns high predicted association probabilities across most indications, with six of eight cases scoring *≥* 0.9, whereas DREAMwalk and K-Paths each exceed this threshold in two cases. Baseline scores also vary substantially across entries, while CAREPath remains high in most disease areas and drug classes. For example, food allergy is a sparsely connected node in the KG, with few associated genes in its local neighborhood. Despite this limited local evidence, CAREPath assigns a high score to this pair by leveraging the shared IgE-neutralizing mechanism with omalizumab’s previously approved allergic indications such as asthma and chronic urticaria [50].

**Table 2.** Predicted probabilities for drug–disease associations corresponding to newly approved indications (2024–2025). Higher values indicate stronger predicted associations. CAREPath achieved consistently high probabilities in six of eight cases.

| New Approved Indication | Drug | CAREPath | DREAMwalk | K-Paths |
| --- | --- | --- | --- | --- |
| Diabetic Retinopathy | Ranibizumab | **0.919** | 0.028 | 0.151 |
| Urothelial Carcinoma | Nivolumab | 0.902 | 0.382 | **0.903** |
| Bipolar I Disorder | Iloperidone | 0.915 | **0.979** | 0.878 |
| Pediatric HeFH | Alirocumab | **0.913** | **0.913** | 0.285 |
| Food Allergy | Omalizumab | **0.913** | 0.859 | 0.806 |
| Ph+ ALL | Ponatinib | 0.917 | 0.510 | **0.936** |
| Atopic Dermatitis | Roflumilast | 0.114 | 0.075 | **0.173** |
| Severe Asthma | Benralizumab | 0.679 | **0.848** | 0.757 |
| # of strong prediction ($\geq 0.9$) | | 6 | 2 | 2 |

## Conclusion

We presented CAREPath, a framework for drug repurposing that combines constrained short disease–gene–drug paths with biomedical language model encoding and mechanism-context from local gene neighborhoods. This design yields accurate predictions with mechanistically interpretable rationales. Across five heterogeneous BKGs, CAREPath achieves the best overall AUPRC, including under disease cold-start settings. Analyses show that semantic path encoding drives the largest gains, while mechanism-context improves robustness for sparsely annotated entities and alignment with GO-BP annotations relative to the DFS-only variant. Evaluation using newly approved FDA indications (2024–2025) suggests that CAREPath assigns high scores to many later approved associations. These results point to a broader observation: deeper multi-hop traversal does not necessarily improve mechanistic reasoning, as longer paths route through hub genes and dilute pair-specific signal. CAREPath shows that constraining path length while recovering mechanism-context balances accuracy and interpretability.

Although CAREPath achieves strong and consistent performance across diverse BKGs, several limitations remain that motivate future extensions. Its current design intentionally relies on short gene-mediated paths and mechanism-context augmentation to balance interpretability, scalability, and robustness. While this design generalizes well across datasets, its performance can still depend on the completeness of the underlying graph, particularly for drugs or diseases with limited gene annotations. Future work can further strengthen CAREPath by incorporating complementary information sources, such as chemical structure, transcriptomic response, or clinical knowledge, to enrich the available mechanistic signal beyond the fixed BKG. In addition, although the short-path constraint effectively reduces hub-driven noise and supports interpretable reasoning, some regulatory mechanisms may involve longer gene cascades that extend beyond the current traversal budget. Extending CAREPath with biologically guided path filtering or adaptive path selection could capture such mechanisms while preserving the noise-control advantages of constrained traversal. These extensions would broaden the framework toward multi-modal and context-adaptive drug repurposing while maintaining its central goal of scalable, mechanism-aware prediction.

### Key Points

- CAREPath is a KG–LLM framework for drug repurposing, combining constrained short disease–gene–drug paths (DFS) with mechanism-context embeddings (BFS).
- Across five biomedical knowledge graphs and 18 baselines, CAREPath achieves the best overall AUPRC and remains robust under sparse and abundant gene neighborhoods.
- Case studies show that CAREPath recovers clinically meaningful regulatory pathways with interpretable rationales, and it assigns high scores to recently FDA-approved indications.

## Data availability

The source code of CAREPath is available at https://github.com/hamppy-song/CAREPath. All datasets analysed in this study are publicly available: MSI [16], PrimeKG [17], Hetionet [10], SuppKG [18], and KEGG50k [19].

## Footnotes

1 Retrieved from https://www.drugs.com/new-indications.html.

