## Supplementary Material for "CAREPath: Semantic Context-Aware Reasoning Paths with Mechanism-Augmented Embeddings for Drug Repurposing"

### Supplementary Methods

#### S1. Baselines

We evaluated our approach against a diverse set of baseline models to benchmark its performance. These models are categorized into four groups based on their underlying principles.

- **GNN-based models**: These models learn node representations by aggregating information from a node's local neighborhood.
  - **GraphSAGE** [1]: An inductive GNN model designed to scale to large graphs. Instead of using all neighbors, it samples a fixed number of neighbors and aggregates their features using functions like mean or max-pooling. This allows it to generalize to unseen nodes.
  - **CompGCN** [2]: A relational GNN that jointly embeds entities and relations by incorporating relation composition operators into the message-passing process. CompGCN updates node representations by explicitly modeling edge types, making it suitable for multi-relational knowledge graphs.
  - **GIN** [3]: A powerful GNN architecture that is theoretically as expressive as the Weisfeiler-Lehman graph isomorphism test. GIN focuses on learning a strong aggregation function that can distinguish between different graph structures, making it highly effective for graph classification tasks.
  - **GAT** [4]: This model introduces an attention mechanism into the GNN framework. Instead of treating all neighbors equally, GAT learns a weight for each neighbor's contribution to the central node's representation. This allows the model to selectively focus on the most relevant neighbors, which is particularly useful for heterogeneous graphs.
- **Triplet-based models**: These models, also known as Knowledge Graph Embedding (KGE) models, learn to represent entities (like drugs and diseases) and relations as vectors in a continuous vector space.
  - **TransE** [5]: A simple but influential model that represents relations as a translation vector. It assumes that for a valid triplet $(h, r, t)$, the embedding of the head entity $h$ plus the relation vector $r$ should be close to the tail entity $t$ (i.e., $h + r \approx t$).
  - **TransR** [6]: An extension of TransE that addresses its limitations by projecting entities into a relation-specific vector space before performing the translation. This allows the model to better handle complex and varied relations.
  - **RotatE** [7]: This model represents relations as a rotation from the head entity to the tail entity in a complex vector space. This rotational mechanism is highly effective at capturing various relation patterns, including symmetry, anti-symmetry, and composition.
  - **ComplEx** [8]: An embedding model that represents entities and relations using complex-valued vectors. It uses a bilinear scoring function to capture asymmetric relationships, which are common in knowledge graphs.
  - **RESCAL** [9]: This is a tensor factorization-based model that represents each relation as a full matrix. It learns a bilinear interaction between entity embeddings to predict the plausibility of a triplet, allowing it to model rich, complex relationships but with higher computational cost.
- **Pretrained biomedical language models**: These models are a different class of baselines that leverage the vast amount of biomedical text data available to learn rich, contextualized representations of biomedical entities.
  - **BioBERT** [10]: A BERT-based language model further pre-trained on biomedical literature, including PubMed abstracts and PubMed Central full-text articles, to better capture domain-specific terminology and semantics.
  - **PubMedBERT** [11]: A domain-specific BERT model trained from scratch exclusively on PubMed abstracts and full-text articles, without relying on general-domain pretraining, enabling representations tailored to biomedical text.

- **BioLinkBERT** [12]: A biomedical language model pre-trained with explicit modeling of document-level and entity-level links, designed to capture relational and contextual information across biomedical texts.
- **SapBERT** [13]: This model builds upon a biomedical language model (like PubMedBERT) but is further fine-tuned using a metric learning framework on massive biomedical ontologies (e.g., UMLS). Its primary goal is to align the representation space of synonymous or related entities, making it particularly powerful for entity linking tasks.

- **Recent Representative models**: These models capture information by exploring multi-hop paths and relational contexts within the knowledge graph, often through random walks.
  - **Node2Vec** [14]: A seminal model for learning node embeddings from random walks. It generates a series of random walks from each node and then uses an algorithm similar to Word2Vec to learn embeddings that capture both local neighborhood structure and global network position.
  - **DrugRep-KG** [15]: A framework that constructs a drug-disease knowledge graph and uses a Word2Vec-like approach on the paths to create numerical vectors for entities and relationships. It specifically focuses on simplifying the relationships between drugs and diseases to better capture potential associations for repurposing.
  - **DREAMwalk** [16]: This model extends the concept of random walks by incorporating semantic information, like ATC codes, to guide the walk. When a walker lands on a drug or disease node, it can teleport to other semantically similar nodes. This process generates paths that are both biologically and semantically meaningful, leading to more informative embeddings.
  - **FuseLinker** [17]: A GNN-based link prediction framework for biomedical knowledge graphs that integrates graph structural information with external textual and domain knowledge embeddings. FuseLinker initializes node representations by fusing pre-trained text embeddings obtained from large language models with domain knowledge embeddings derived from biomedical ontologies, and subsequently refines them using a relational GNN encoder for link prediction tasks.
  - **K-Paths** [18]: A path-based reasoning model that performs inference over sets of multi-hop paths between drug and disease entities in a knowledge graph. K-Paths explicitly enumerates and encodes informative $k$-hop graph paths and applies neural reasoning over these paths to support drug repurposing and drug–drug interaction prediction.

#### S2. Similarity Definitions and Pooling Strategy

##### Mechanism Context Embeddings from One-hop Gene Neighborhoods

For each entity, we first build an entity-centric gene neighborhood context from the biomedical KG $\mathcal{G} = (\mathcal{V}, \mathcal{E})$. Let $\mathcal{V}_{\text{gene}}$ denote gene/protein nodes. For a disease $s$ and a drug $d$, we collect their one-hop gene neighborhoods

$$\mathcal{N}_g(s) = \{g \in \mathcal{V}_{\text{gene}} \mid (s, g) \in \mathcal{E} \text{ or } (g, s) \in \mathcal{E}\}, \qquad \mathcal{N}_g(d) = \{g \in \mathcal{V}_{\text{gene}} \mid (d, g) \in \mathcal{E} \text{ or } (g, d) \in \mathcal{E}\}.$$

To avoid leakage from drug–disease co-neighborhoods, we retain only gene/protein-type neighbors when constructing context sentences. Each neighbor is mapped to a human-readable name and converted into a sentence:

$$t(d, g) = \texttt{Drug } d \texttt{ is connected to } g\texttt{.}, \qquad t(s, g) = \texttt{Disease } s \texttt{ is connected to } g\texttt{.}$$

Let $T(d) = \{t(d, g) : g \in \mathcal{N}_g(d)\}$ and $T(s) = \{t(s, g) : g \in \mathcal{N}_g(s)\}$. We encode each sentence using a frozen biomedical encoder $\phi(\cdot)$ (BioLinkBERT) and obtain initial mechanism context embeddings by mean pooling:

$$\mathbf{Z}^{\text{drug}}_{\text{gene}}(d) = \frac{1}{|T(d)|} \sum_{t \in T(d)} \phi(t), \qquad \mathbf{Z}^{\text{dis}}_{\text{gene}}(s) = \frac{1}{|T(s)|} \sum_{t \in T(s)} \phi(t).$$

If $\mathcal{N}_g(v) = \emptyset$, we use a single fallback sentence to ensure $|T(v)| = 1$.

##### Drug Similarity via ATC Grouping

To obtain pharmacologically related drugs, we use ATC-code grouping. Let $\text{ATC}_L(d)$ denote the first $L$ characters of the ATC code of drug $d$ (in our implementation, $L = 3$ was used to build the teleport/similarity graph, and $L = 4$ was used for context pooling). We define the drug similarity neighbor set as

$$\mathcal{K}^{\text{drug}}(d) = \{d' \in \mathcal{V}_{\text{drug}} \setminus \{d\} \mid \text{ATC}_L(d') = \text{ATC}_L(d)\}.$$

The ATC-based pooled embedding is computed by simple mean pooling over the embeddings of drugs in the same ATC group:

$$\mathbf{Z}_{\text{sim}}^{\text{drug}}(d) = \begin{cases} \frac{1}{|\mathcal{K}^{\text{drug}}(d)|} \sum_{d' \in \mathcal{K}^{\text{drug}}(d)} \mathbf{Z}_{\text{gene}}^{\text{drug}}(d'), & |\mathcal{K}^{\text{drug}}(d)| > 0, \\ \mathbf{0}, & \text{otherwise.} \end{cases}$$

Finally, we apply residual mixing:

$$\mathbf{Z}_{\text{ctx}}^{\text{drug}}(d) = \beta\, \mathbf{Z}_{\text{gene}}^{\text{drug}}(d) + (1-\beta)\, \mathbf{Z}_{\text{sim}}^{\text{drug}}(d),$$

where $\beta \in [0,1]$ (we used $\beta = 0.5$ unless otherwise noted).

##### Disease Similarity via Weighted Gene-neighborhood kNN

For diseases, we define similarity based on overlap of one-hop gene neighborhoods, with downweighting of hub genes. Let $\mathcal{S}$ be the set of disease nodes. We construct a sparse weighted gene vector $\mathbf{x}_s \in \mathbb{R}^{|\mathcal{G}|}$ for each disease $s \in \mathcal{S}$:

$$x_{s,g} = \begin{cases} \text{idf}(g), & g \in \mathcal{N}_g(s), \\ 0, & \text{otherwise,} \end{cases}$$

where $\text{idf}(g)$ is an inverse-document-frequency weight computed across diseases:

$$\text{idf}(g) = \log\left(\frac{|\mathcal{S}|+1}{\text{df}(g)+1}\right) + 1, \qquad \text{df}(g) = |\{s \in \mathcal{S} : g \in \mathcal{N}_g(s)\}|.$$

We then row-normalize these vectors and compute cosine similarity. Let

$$\tilde{\mathbf{x}}_s = \frac{\mathbf{x}_s}{\|\mathbf{x}_s\|_2 + \epsilon}.$$

We define the similar disease set $\mathcal{K}^{\text{disease}}(s)$ as the top-$k$ nearest neighbors of $s$ in cosine distance over $\tilde{\mathbf{x}}_s$ (excluding $s$ itself). Using cosine distance $\text{dist}(\cdot,\cdot)$, similarity is $\text{sim}(s,s') = 1 - \text{dist}(\tilde{\mathbf{x}}_s, \tilde{\mathbf{x}}_{s'})$. We pool neighbor embeddings with similarity-weighted mean:

$$\mathbf{Z}_{\text{sim}}^{\text{dis}}(s) = \begin{cases} \sum_{s' \in \mathcal{K}^{\text{disease}}(s)} w_{s,s'}\, \mathbf{Z}_{\text{gene}}^{\text{dis}}(s'), & |\mathcal{K}^{\text{disease}}(s)| > 0, \\ \mathbf{0}, & \text{otherwise,} \end{cases} \qquad w_{s,s'} = \frac{\text{sim}(s,s')}{\sum_{u \in \mathcal{K}^{\text{disease}}(s)} \text{sim}(s,u) + \epsilon}.$$

Finally, we apply residual mixing:

$$\mathbf{Z}_{\text{ctx}}^{\text{dis}}(s) = \alpha\, \mathbf{Z}_{\text{gene}}^{\text{dis}}(s) + (1-\alpha)\, \mathbf{Z}_{\text{sim}}^{\text{dis}}(s),$$

where $\alpha \in [0,1]$ (we used $\alpha = 0.5$) and $k = 10$ in our implementation.

##### Notes on Robustness Controls

To mitigate extreme hub effects, we cap the maximum number of one-hop gene neighbors used to build the disease gene vectors (500 in our implementation) and optionally subsample overly large neighborhoods for sentence construction. All similarity computations are performed using the KG neighborhood structure only; drug–disease edges are not used when forming entity contexts to prevent label leakage.

#### S3. Dataset

Table S1: Summary statistics of the knowledge graphs and the drug–disease association datasets.

| Datasets | # of nodes | # of edges | Training | Validation | Test |
|---|---|---|---|---|---|
| MSI | 41,941 | 478,728 | 9,488 | 1,186 | 1,186 |
| PrimeKG | 88,357 | 1,340,022 | 13,969 | 1,746 | 1,746 |
| Hetionet | 19,918 | 1,114,451 | 1,194 | 149 | 149 |
| SuppKG | 26,615 | 147,038 | 33,360 | 4,170 | 4,170 |
| KEGG50k | 16,201 | 63,080 | 8,988 | 1,124 | 1,124 |

Table S1 summarizes the datasets used in this study. For each knowledge graph, we report the total number of nodes and edges in the graph, together with the number of labeled drug–disease pairs used for supervised association prediction. The labeled pairs were split into training, validation, and test sets using an 8:1:1 ratio. Note that the graph statistics (# of nodes and # of edges) describe the full knowledge graph structure, whereas the split sizes correspond to the labeled drug–disease pairs only.

#### MSI network

We used the Multiscale Interactome (MSI) network[1], a disease–treatment explanation graph that integrates drugs, diseases, human proteins, and a hierarchy of biological functions to support mechanistic interpretation of therapeutic effects [? ]. We downloaded the original MSI graph and filtered it to retain only the node types required for our mechanistic setting: drugs, diseases, genes/proteins, and biological functions (GO terms). We further restricted the graph to five relation types: drug–gene, disease–gene, gene–gene, gene–biological function, and biological function–biological function. After filtering, the resulting MSI subgraph contains 41,941 nodes and 478,728 edges. Labeled drug–disease pairs were used for supervised prediction and were split into training, validation, and test sets (8:1:1).

#### PrimeKG

PrimeKG is a precision-medicine knowledge graph[2] that consolidates diverse biomedical resources to connect diseases with therapeutic, molecular, and phenotypic evidence across multiple biological scales [19]. Starting from the original PrimeKG release, we filtered the graph to retain only node types relevant to our setting: genes/proteins, drugs, diseases, biological process, anatomy, molecular function, cellular component, and pathway. Our processed PrimeKG graph contains 1,340,022 edges and 88,357 nodes.

#### Hetionet

Hetionet is a heterogeneous biomedical network curated for systematic drug repurposing via integration of public data sources spanning compounds, diseases, genes, pathways, and functional ontologies [20]. We downloaded the original Hetionet release and filtered it to match the entity scope of our framework by retaining only drug, disease, and gene nodes and removing all other node and relation types. We further extracted drug–disease association edges to construct the labeled prediction dataset, which was split into training/validation/test sets (8:1:1). After filtering, the resulting Hetionet subgraph contains 19,918 nodes and 1,114,451 edges.

#### SuppKG

SuppKG is a literature-derived knowledge graph designed to represent dietary supplement concepts that are under-covered in standard biomedical terminologies. It is constructed from PubMed abstracts using an extended SemRep pipeline enriched with a supplement-specific terminology, with additional relation filtering to improve precision [21]. Entities in SuppKG are annotated with UMLS semantic type (semtype) codes. To align SuppKG with our mechanistic setting, we constructed a task-aligned subgraph by retaining only drug-, disease-, and gene/protein-related concepts based on selected semtypes: `phsu` and `orch` (drug), `dsyn` (disease), and `gngm`, `aapp`, and `enzy` (gene/protein). We then filtered edges to keep only those whose endpoints both belonged to the retained node sets, yielding a DDG-only subgraph. This filtered SuppKG subgraph contains 26,615 nodes and 147,038 edges. For evaluation, labeled drug–disease pairs were split into training/validation/test sets in an 8:1:1 ratio using stratified sampling.

#### KEGG50k

KEGG is a curated biological systems resource that represents molecular networks with an emphasis on manually constructed pathway maps and functional organization via orthology [22]. Based on KEGG, KEGG50k has been released as a ready-to-download benchmarking biomedical knowledge graph for evaluating knowledge-graph embedding models, originally curated to support drug–target interaction prediction while preserving pathway-structured biological context (e.g., drug–target, disease–gene, and pathway–gene links).[3] We downloaded this KEGG50k release and used it as provided. Our processed KEGG50k graph contains 16,201 nodes and 63,080 edges. For supervised evaluation, we used labeled drug_target_disease pairs and split them into training, validation, test sets (8:1:1).

#### Negative Sampling

To generate negative samples, we randomly sampled drug–disease pairs that exist as nodes in the graph but are not connected by any known association edge. For MSI, Hetionet, SuppKG, and KEGG50k, we used a 1:1 positive-to-negative ratio via random sampling. For PrimeKG, we used a slightly higher 1:1.2 ratio (7,461 positives and 10,000 negatives) to reflect its broader disease coverage.

[1]https://github.com/snap-stanford/multiscale-interactome
[2]https://github.com/mims-harvard/PrimeKG
[3]`https://figshare.com/s/bbfc7b82d17e0b8b6a43`

### S4. Experimental Setup

#### S4.1 Hyperparameters of CAREPath (Proposed Model)

Hyperparameters were optimized via grid search separately for each knowledge graph, and the selected configurations are summarized in Table S2. In particular, Table S1 reports the best-performing XGBoost hyperparameters used across all experiments.

Table S2: Selected hyperparameters for XGBoost (and logistic regression meta-model when applicable) across knowledge graphs.

| Hyperparameter | MSI | PrimeKG | Hetionet | SuppKG | KEGG50k |
|---|---|---|---|---|---|
| learning_rate | 0.0221 | 0.0648 | 0.0212 | 0.1527 | 0.0639 |
| n_estimators | 1597 | 1209 | 1559 | 859 | 932 |
| max_depth | 7 | 5 | 5 | 6 | 10 |
| min_child_weight | 13 | 16 | 1 | 4 | 3 |
| subsample | 0.6540 | 0.7221 | 0.6110 | 0.9388 | 0.8335 |
| colsample_bytree | 0.8105 | 0.5879 | 0.6326 | 0.9006 | 0.8822 |
| gamma | 0.3203 | 0.6465 | 1.05e-04 | 1.0170 | 0.5810 |
| reg_alpha | 6.09e-07 | 0.3319 | 1.49e-06 | 0.0038 | 0 |
| reg_lambda | 0.6138 | 13.2332 | 0.0015 | 0.5607 | 4.2877 |
| scale_pos_weight | 1.0324 | 1.3188 | 1.8299 | 1.000 | 1.000 |

For all datasets, we use the following hyperparameters for constructing CAREPath embeddings.

- **Node2Vec (node embeddings).** Embedding dimension = 128, walk length = 10, number of walks per node = 100. The skip-gram model is trained with context window size = 10 (min_count = 1).
- **Path extraction.** We enumerate disease–gene–drug paths up to 3 hops (cutoff = 3), and limit the number of intermediate gene/protein nodes in a path to at most 2 (max_genes = 2). If no path exists, we use a fallback prompt with empty gene evidence.
- **Drug-side teleportation (ATC-guided).** When no fixed path is found for a given pair, we apply ATC-group-based teleportation with probability 0.3 to sample alternative drug nodes within the same ATC prefix group.
- **BioLinkBERT path encoder.** We encode each path as an NLI-style prompt (Premise/Hypothesis) and extract the final-layer `[CLS]` representation. For a disease–drug pair, multiple path embeddings are aggregated by element-wise max pooling.
- **Mechanism context embedding.** For each drug/disease node, we build context texts from its 1-hop gene/protein neighbors only (excluding drug–disease neighbors), and obtain an entity-level mechanism embedding by mean pooling the CLS embeddings of all available context sentences.
- **Neighbor pooling for mechanism embeddings.** For drugs, we pool within the same ATC group and mix it with the original mechanism embedding using residual mixing with $\alpha = 0.5$. For diseases, we compute sparse weighted gene vectors (IDF-weighted; with at most 500 sampled 1-hop genes per node), retrieve top-$k$ neighbors using cosine similarity ($k = 10$), and apply the same residual mixing with $\beta = 0.5$.

#### S4.2 Other Experimental Details

All experiments were conducted on a single NVIDIA RTX 3090 GPU (24GB) running Ubuntu 18.04. We report benchmark results using 5-fold cross-validation with a fixed random seed (42), summarized as mean±std over folds. We consider three splitting protocols: (i) **Random split**, which applies stratified 5-fold cross-validation (`StratifiedKFold`) over labeled drug–disease pairs to preserve the class ratio in each fold; (ii) **Disease split**, which uses group-based cross-validation (`GroupKFold`) grouping by disease so that all pairs sharing the same disease appear in the same fold (i.e., test diseases are unseen during training); and (iii) **Drug split**, which analogously uses `GroupKFold` grouping by drug so that test drugs are unseen during training. In all settings, each fold trains on 4 folds and evaluates on the held-out fold.

#### S4.3 Baseline models

**Graph-based models** All graph-only baselines use a message-passing encoder and a pairwise scoring head that concatenates the final drug and disease node embeddings and applies a linear layer with sigmoid output. For GraphSAGE, GAT, and GIN, we use a 2-layer encoder with hidden dimension 64 and train with Adam for 50 epochs. For GAT, we use 2 attention heads in the first layer with dropout 0.2. For CompGCN+DistMult, we use a 2-layer CompGCN encoder (hidden dimension 128; composition `mult`; dropout 0.1) with a DistMult decoder, trained with Adam for 50 epochs (learning rate $2 \times 10^{-3}$, weight decay $1 \times 10^{-5}$; batch size 4096).

**KGE-based models** We evaluated five knowledge graph embedding (KGE) baselines—TransE, TransR, RotatE, ComplEx, and RESCAL—under the same splitting and evaluation protocol as our main benchmarks. For all KGE models, we trained embeddings using Adam with learning rate 0.01 for 100 epochs and used an embedding dimension of 128 (with relation dimension 64 for TransR). We used a standard margin-based ranking objective with negative samples generated by random corruption during training. To obtain supervised drug–disease predictions from the learned embeddings, we constructed a pair feature from the drug and disease embeddings (element-wise absolute difference) and trained a downstream ensemble classifier. For this step, we reused the same XGBoost stacking configuration as our main pipeline for each dataset (the downstream XGBoost ensemble hyperparameters were fixed per dataset, and the same dataset-specific configuration in Table S2 was used consistently across all KGE variants.), ensuring a consistent and fair comparison across KGE baselines.

**LLM-based models** We evaluate LLM-based baselines that encode each drug–disease pair into a fixed-length vector using a frozen biomedical language model, without any KG message passing or path-derived features. We use four encoders (BioBERT, BioLinkBERT, PubMedBERT, and SapBERT) under an identical protocol. For each pair $(d, s)$, we map entity IDs to human-readable names using the same ID-to-name dictionary as in our main pipeline, and construct an input prompt with a fixed maximum sequence length (128 tokens) using padding/truncation. We extract the final-layer `[CLS]` embedding as the pair representation and keep the encoder frozen during training.

The prompt follows the same format used in our main model:

> Premise: $s$ involves genes $\{g_1, \ldots, g_k\}$.
> Hypothesis: $d$ can be repurposed to treat $s$.
> Label:

For classification, we standardize embeddings with `StandardScaler` and train the same downstream XGBoost stacking ensemble used elsewhere for each dataset (Table S2).

**Recent Representative Models**

- **Node2Vec**: For each knowledge graph, we learn unsupervised node representations using Node2Vec. We construct a graph from the edge list and generate biased random-walk sequences with embedding dimension 128, walk length 10, and 100 walks per node. The resulting walk corpus is used to train a skip-gram Word2Vec model with context window size 10 (minimum count = 1), producing a 128-dimensional embedding for every node. For each disease–drug pair $(s, d)$, we form the pair feature by concatenating the corresponding node embeddings, i.e., $[\mathbf{z}_s; \mathbf{z}_d]$, and use it as input to downstream prediction models.

- **Drugrep-KG (2023)**: For the DrugRep-KG baseline, we trained a CBOW Word2Vec model on KG-derived triplet sentences with embedding dimension 128, context window size 5, minimum count 1, and 10 training epochs. Drug–disease pair features were formed by concatenating the learned drug and disease embeddings and were standardized before classification. For each dataset, we reused the same dataset-specific downstream classifier configuration as in the main pipeline to ensure a consistent comparison.

- **DREAMwalk (2023)**: For the DREAMwalk baseline, we followed the hyperparameter settings reported in the original DREAMwalk paper without additional tuning. For each dataset, we used the same training and evaluation protocol as in our main pipeline to ensure a consistent comparison.

- **FuseLinker (2024)**: For the FuseLinker baseline, we used the Text Embedding Module together with the GNN Model Module, while excluding the Domain Knowledge Embedding Module (the UMLS/MetaMap-based semantic-type mapping and ontology embedding component). This usage is consistent with recent work [23] that adopts FuseLinker as a baseline/competitor. All hyperparameters were tuned separately for each dataset using a validation-set grid search following the original experimental setup.

- **K-Paths (2025)**: We used the LLM-based reasoning track of K-Paths (i.e., without the GNN-based subgraph prediction track). We retrieved $K = 10$ paths per query and capped the maximum path length at 3, then provided the LLM with the resulting textualized paths. We used Tx-Gemma-9B-Chat for path-based inference. All hyperparameters were tuned separately for each dataset via a validation-set grid search following the original experimental protocol while keeping the model backbone fixed.

### S5. Detailed Results

#### Random Split

Table S3: Performance comparison on MSI (Random Split).

| Methods | Models | AUROC | AUPRC | Accuracy | F1 |
|---|---|---|---|---|---|
| Graph-based | GraphSAGE [1] | 0.737±0.004 | 0.711±0.004 | 0.696±0.005 | 0.694±0.006 |
| | CompGCN [2] | 0.641±0.011 | 0.633±0.012 | 0.607±0.010 | 0.553±0.021 |
| | GIN [3] | 0.689±0.073 | 0.694±0.071 | 0.638±0.069 | 0.640±0.036 |
| | GAT [4] | 0.776±0.002 | 0.763±0.004 | 0.715±0.008 | 0.732±0.007 |
| KGE-based | TransE [5] | 0.748±0.009 | 0.771±0.008 | 0.685±0.009 | 0.668±0.011 |
| | TransR [6] | 0.661±0.014 | 0.698±0.018 | 0.613±0.015 | 0.592±0.006 |
| | RotatE [7] | 0.583±0.015 | 0.602±0.017 | 0.530±0.016 | 0.632±0.006 |
| | ComplEx [8] | 0.744±0.018 | 0.763±0.018 | 0.683±0.016 | 0.659±0.019 |
| | RESCAL [9] | 0.678±0.006 | 0.700±0.009 | 0.627±0.006 | 0.609±0.013 |
| LLM-based | BioBERT [10] | 0.833±0.008 | 0.832±0.009 | 0.768±0.008 | 0.768±0.008 |
| | BioLinkBERT [12] | 0.888±0.006 | 0.890±0.007 | 0.813±0.007 | 0.812±0.006 |
| | PubMedBERT [11] | 0.902±0.005 | 0.904±0.005 | 0.830±0.005 | 0.830±0.005 |
| | SapBERT [13] | 0.874±0.003 | 0.877±0.002 | 0.800±0.008 | 0.798±0.006 |
| Path-based | Node2Vec [14] | 0.890±0.003 | 0.899±0.003 | 0.815±0.002 | 0.814±0.002 |
| | DrugRep-KG [15] | 0.890±0.003 | 0.899±0.006 | 0.814±0.003 | 0.812±0.004 |
| | DREAMwalk [16] | 0.900±0.004 | 0.908±0.005 | 0.826±0.004 | 0.825±0.004 |
| | FuseLinker [17] | 0.902±0.003 | 0.896±0.025 | 0.853±0.034 | **0.861±0.021** |
| | K-Paths [18] | 0.910±0.004 | 0.913±0.023 | 0.831±0.014 | 0.824±0.016 |
| | CAREPath | **0.932±0.003** | **0.936±0.004** | **0.861±0.001** | **0.861±0.003** |

Table S4: Performance comparison on PrimeKG (Random Split).

| Methods | Models | AUROC | AUPRC | Accuracy | F1 |
|---|---|---|---|---|---|
| Graph-based | GraphSAGE [1] | 0.976±0.001 | 0.960±0.005 | 0.923±0.003 | 0.914±0.003 |
| | CompGCN [2] | 0.947±0.002 | 0.927±0.006 | 0.871±0.012 | 0.855±0.008 |
| | GIN [3] | 0.859±0.049 | 0.816±0.058 | 0.766±0.057 | 0.683±0.103 |
| | GAT [4] | 0.978±0.003 | 0.964±0.008 | 0.931±0.005 | 0.922±0.005 |
| KGE-based | TransE [5] | 0.938±0.010 | 0.942±0.008 | 0.888±0.012 | 0.861±0.015 |
| | TransR [6] | 0.818±0.043 | 0.819±0.052 | 0.742±0.056 | 0.714±0.039 |
| | RotatE [7] | 0.768±0.017 | 0.721±0.020 | 0.569±0.020 | 0.656±0.010 |
| | ComplEx [8] | 0.967±0.001 | 0.966±0.001 | 0.906±0.003 | 0.880±0.005 |
| | RESCAL [9] | 0.941±0.003 | 0.939±0.001 | 0.863±0.002 | 0.818±0.004 |
| LLM-based | BioBERT [10] | 0.971±0.002 | 0.962±0.002 | 0.910±0.004 | 0.896±0.005 |
| | BioLinkBERT [12] | 0.959±0.002 | 0.943±0.004 | 0.889±0.003 | 0.871±0.003 |
| | PubMedBERT [11] | 0.964±0.002 | 0.954±0.004 | 0.897±0.005 | 0.879±0.006 |
| | SapBERT [13] | 0.944±0.004 | 0.928±0.006 | 0.867±0.008 | 0.844±0.010 |
| Path-based | Node2Vec [14] | **0.989±0.001** | **0.987±0.002** | **0.955±0.002** | **0.948±0.002** |
| | DrugRep-KG [15] | 0.768±0.006 | 0.816±0.021 | 0.717±0.005 | 0.774±0.007 |
| | DREAMwalk [16] | 0.988±0.002 | 0.985±0.002 | 0.953±0.003 | 0.945±0.003 |
| | FuseLinker [17] | 0.920±0.005 | 0.913±0.002 | 0.873±0.032 | 0.869±0.009 |
| | K-Paths [18] | 0.737±0.034 | 0.746±0.0021 | 0.710±0.008 | 0.703±0.050 |
| | CAREPath | **0.989±0.001** | 0.985±0.001 | 0.949±0.003 | 0.940±0.003 |

Table S5: Performance comparison on Hetionet (Random Split).

| Methods | Models | AUROC | AUPRC | Accuracy | F1 |
|---|---|---|---|---|---|
| Graph-based | GraphSAGE [1] | 0.855±0.016 | 0.821±0.016 | 0.790±0.018 | 0.799±0.019 |
| | CompGCN [2] | 0.784±0.043 | 0.783±0.043 | 0.690±0.037 | 0.674±0.107 |
| | GIN [3] | 0.525±0.051 | 0.515±0.031 | 0.525±0.049 | 0.668±0.002 |
| | GAT [4] | 0.878±0.016 | 0.843±0.033 | 0.823±0.015 | 0.827±0.019 |
| KGE-based | TransE [5] | 0.598±0.031 | 0.605±0.018 | 0.502±0.018 | 0.661±0.014 |
| | TransR [6] | 0.698±0.051 | 0.715±0.060 | 0.609±0.045 | 0.671±0.018 |
| | RotatE [7] | 0.684±0.034 | 0.696±0.014 | 0.605±0.046 | 0.668±0.035 |
| | ComplEx [8] | 0.789±0.026 | 0.784±0.036 | 0.712±0.028 | 0.736±0.021 |
| | RESCAL [9] | 0.716±0.033 | 0.711±0.048 | 0.656±0.026 | 0.650±0.038 |
| LLM-based | BioBERT [10] | 0.896±0.004 | 0.902±0.003 | 0.819±0.006 | 0.820±0.006 |
| | BioLinkBERT [12] | 0.952±0.018 | 0.952±0.014 | 0.885±0.024 | 0.884±0.024 |
| | PubMedBERT [11] | 0.953±0.011 | 0.953±0.013 | 0.883±0.012 | 0.883±0.013 |
| | SapBERT [13] | 0.930±0.015 | 0.935±0.013 | 0.855±0.013 | 0.854±0.016 |
| Path-based | Node2Vec [14] | 0.940±0.018 | 0.935±0.022 | 0.866±0.023 | 0.868±0.023 |
| | DrugRep-KG [15] | 0.962±0.002 | **0.982±0.001** | **0.904±0.002** | **0.933±0.001** |
| | DREAMwalk [16] | 0.840±0.017 | 0.936±0.017 | 0.869±0.013 | 0.871±0.013 |
| | FuseLinker [17] | 0.937±0.003 | 0.952±0.002 | 0.846±0.030 | 0.878±0.029 |
| | K-Paths [18] | 0.945±0.004 | 0.956±0.002 | 0.890±0.010 | 0.893±0.012 |
| | CAREPath | **0.969±0.008** | 0.971±0.008 | 0.900±0.016 | 0.901±0.017 |

Table S6: Performance comparison on SuppKG (Random Split).

| Methods | Models | AUROC | AUPRC | Accuracy | F1 |
|---|---|---|---|---|---|
| Graph-based | GraphSAGE [1] | 0.915±0.005 | 0.951±0.004 | 0.863±0.004 | 0.903±0.003 |
| | CompGCN [2] | 0.944±0.002 | 0.975±0.001 | 0.881±0.003 | 0.915±0.003 |
| | GIN [3] | 0.872±0.127 | 0.923±0.088 | 0.836±0.068 | 0.887±0.033 |
| | GAT [4] | 0.823±0.021 | 0.848±0.027 | 0.838±0.005 | 0.889±0.003 |
| KGE-based | TransE [5] | 0.911±0.001 | 0.924±0.001 | 0.837±0.002 | 0.827±0.002 |
| | TransR [6] | 0.763±0.023 | 0.767±0.025 | 0.648±0.023 | 0.716±0.011 |
| | RotatE [7] | 0.727±0.006 | 0.732±0.005 | 0.521±0.003 | 0.673±0.001 |
| | ComplEx [8] | 0.907±0.005 | 0.911±0.005 | 0.786±0.017 | 0.813±0.010 |
| | RESCAL [9] | 0.873±0.007 | 0.871±0.007 | 0.792±0.007 | 0.783±0.007 |
| LLM-based | BioBERT [10] | 0.924±0.002 | 0.930±0.002 | 0.849±0.004 | 0.850±0.004 |
| | BioLinkBERT [12] | 0.930±0.003 | 0.934±0.003 | 0.852±0.005 | 0.854±0.005 |
| | PubMedBERT [11] | 0.934±0.004 | 0.939±0.004 | 0.857±0.005 | 0.858±0.004 |
| | SapBERT [13] | 0.923±0.004 | 0.929±0.003 | 0.843±0.004 | 0.846±0.003 |
| Path-based | Node2Vec [14] | 0.959±0.003 | **0.981±0.001** | 0.895±0.003 | 0.927±0.002 |
| | DrugRep-KG [15] | 0.912±0.005 | 0.943±0.004 | 0.851±0.003 | 0.870±0.004 |
| | DREAMwalk [16] | 0.961±0.003 | **0.981±0.002** | 0.897±0.004 | **0.928±0.003** |
| | FuseLinker [17] | 0.896±0.003 | 0.917±0.031 | 0.846±0.012 | 0.860±0.005 |
| | K-Paths [18] | 0.871±0.033 | 0.890±0.016 | 0.813±0.005 | 0.833±0.014 |
| | CAREPath | **0.973±0.001** | 0.975±0.001 | **0.918±0.002** | 0.918±0.002 |

Table S7: Performance comparison on KEGG50k (Random Split).

| Methods | Models | AUROC | AUPRC | Accuracy | F1 |
|---|---|---|---|---|---|
| Graph-based | GraphSAGE [1] | 0.958±0.003 | 0.904±0.012 | 0.905±0.005 | 0.851±0.008 |
| | CompGCN [2] | 0.879±0.010 | 0.757±0.026 | 0.656±0.009 | 0.644±0.006 |
| | GIN [3] | 0.963±0.003 | 0.914±0.007 | 0.930±0.006 | 0.890±0.009 |
| | GAT [4] | 0.967±0.001 | 0.925±0.008 | 0.929±0.004 | 0.890±0.005 |
| KGE-based | TransE [5] | 0.839±0.025 | 0.865±0.019 | 0.658±0.037 | 0.728±0.021 |
| | TransR [6] | 0.772±0.016 | 0.807±0.015 | 0.658±0.028 | 0.699±0.013 |
| | RotatE [7] | 0.651±0.002 | 0.666±0.008 | 0.514±0.004 | 0.667±0.001 |
| | ComplEx [8] | 0.925±0.007 | 0.937±0.006 | 0.861±0.003 | 0.847±0.005 |
| | RESCAL [9] | 0.819±0.012 | 0.851±0.009 | 0.745±0.008 | 0.684±0.015 |
| LLM-based | BioBERT [10] | 0.974±0.002 | 0.971±0.002 | 0.923±0.005 | 0.924±0.004 |
| | BioLinkBERT [12] | 0.982±0.001 | 0.982±0.001 | 0.932±0.003 | 0.932±0.003 |
| | PubMedBERT [11] | 0.984±0.002 | <u>0.983±0.001</u> | 0.938±0.006 | 0.938±0.006 |
| | SapBERT [13] | 0.967±0.002 | 0.968±0.002 | 0.905±0.002 | 0.905±0.002 |
| Path-based | Node2Vec [14] | 0.985±0.004 | 0.977±0.004 | 0.959±0.005 | 0.933±0.009 |
| | DrugRep-KG [15] | <u>0.988±0.001</u> | 0.978±0.003 | <u>0.960±0.005</u> | <u>0.948±0.012</u> |
| | DREAMwalk [16] | 0.991±0.001 | <u>0.983±0.003</u> | **0.963±0.003** | 0.940±0.005 |
| | FuseLinker [17] | 0.975±0.002 | 0.977±0.003 | 0.936±0.003 | 0.935±0.001 |
| | K-Paths [18] | 0.683±0.023 | 0.699±0.011 | 0.645±0.010 | 0.610±0.018 |
| | CAREPath | **0.992±0.002** | **0.991±0.002** | 0.958±0.004 | **0.958±0.004** |

##### Disease Split(Disease Cold-start)

Table S8: Performance comparison on MSI (Disease Cold-start Split).

| Methods | Models | AUROC | AUPRC | Accuracy | F1 |
|---|---|---|---|---|---|
| Graph-based | GraphSAGE [1] | 0.589±0.009 | 0.574±0.016 | 0.577±0.016 | 0.548±0.018 |
| | CompGCN [2] | 0.642±0.024 | 0.631±0.028 | 0.608±0.018 | 0.557±0.028 |
| | GIN [3] | 0.655±0.014 | 0.651±0.017 | 0.609±0.020 | 0.539±0.055 |
| | GAT [4] | 0.621±0.018 | 0.588±0.032 | 0.581±0.016 | 0.574±0.036 |
| KGE-based | TransE [5] | 0.576±0.014 | 0.534±0.013 | 0.536±0.010 | 0.530±0.028 |
| | TransR [6] | 0.551±0.023 | 0.579±0.019 | 0.540±0.017 | 0.575±0.044 |
| | RotatE [7] | 0.540±0.018 | 0.559±0.017 | 0.518±0.009 | 0.551±0.067 |
| | ComplEx [8] | 0.560±0.011 | 0.566±0.018 | 0.504±0.020 | 0.483±0.008 |
| | RESCAL [9] | 0.509±0.010 | 0.501±0.030 | 0.511±0.010 | 0.465±0.105 |
| LLM-based | BioBERT [10] | 0.719±0.013 | 0.706±0.021 | 0.649±0.007 | 0.581±0.035 |
| | BioLinkBERT [12] | 0.841±0.012 | 0.849±0.011 | 0.763±0.008 | 0.741±0.019 |
| | PubMedBERT [11] | **0.904±0.011** | <u>0.857±0.014</u> | 0.791±0.021 | 0.741±0.014 |
| | SapBERT [13] | 0.821±0.015 | 0.830±0.019 | 0.750±0.015 | 0.731±0.014 |
| Path-based | Node2Vec [14] | 0.752±0.023 | 0.766±0.007 | 0.664±0.024 | 0.565±0.036 |
| | DrugRep-KG [15] | 0.733±0.012 | 0.740±0.013 | 0.643±0.026 | 0.620±0.011 |
| | DREAMwalk [16] | 0.763±0.008 | 0.769±0.008 | 0.663±0.116 | 0.561±0.021 |
| | FuseLinker [17] | 0.854±0.014 | 0.806±0.013 | 0.766±0.022 | 0.738±0.015 |
| | K-Paths [18] | 0.819±0.012 | 0.795±0.017 | **0.821±0.020** | **0.788±0.018** |
| | CAREPath | <u>0.890±0.014</u> | **0.894±0.016** | <u>0.802±0.011</u> | <u>0.782±0.017</u> |

Table S9: Performance comparison on PrimeKG (Disease Cold-start Split).

| Methods | Models | AUROC | AUPRC | Accuracy | F1 |
|---|---|---|---|---|---|
| Graph-based | GraphSAGE [1] | 0.901±0.013 | 0.853±0.023 | 0.812±0.024 | 0.764±0.040 |
| | CompGCN [2] | 0.924±0.006 | 0.802±0.055 | 0.856±0.014 | 0.830±0.019 |
| | GIN [3] | 0.832±0.048 | 0.515±0.046 | 0.774±0.042 | 0.697±0.078 |
| | GAT [4] | 0.913±0.010 | 0.751±0.028 | 0.837±0.010 | 0.794±0.013 |
| KGE-based | TransE [5] | 0.714±0.029 | 0.649±0.045 | 0.620±0.022 | 0.600±0.034 |
| | TransR [6] | 0.596±0.030 | 0.547±0.047 | 0.569±0.034 | 0.536±0.076 |
| | RotatE [7] | 0.785±0.017 | 0.742±0.013 | 0.661±0.018 | 0.683±0.010 |
| | ComplEx [8] | 0.794±0.022 | 0.782±0.021 | 0.604±0.018 | 0.661±0.036 |
| | RESCAL [9] | 0.612±0.046 | 0.558±0.036 | 0.573±0.021 | 0.473±0.030 |
| LLM-based | BioBERT [10] | 0.888±0.011 | 0.842±0.012 | 0.755±0.025 | 0.645±0.031 |
| | BioLinkBERT [12] | 0.913±0.006 | 0.878±0.015 | 0.809±0.013 | 0.751±0.024 |
| | PubMedBERT [11] | 0.905±0.009 | 0.868±0.021 | 0.792±0.012 | 0.717±0.032 |
| | SapBERT [13] | 0.900±0.008 | 0.871±0.011 | 0.804±0.013 | 0.747±0.019 |
| Path-based | Node2Vec [14] | 0.944±0.012 | 0.931±0.008 | 0.835±0.021 | 0.771±0.030 |
| | DrugRep-KG [15] | 0.682±0.013 | 0.778±0.029 | 0.563±0.038 | 0.520±0.011 |
| | DREAMwalk [16] | 0.941±0.011 | 0.929±0.014 | 0.838±0.025 | 0.777±0.041 |
| | FuseLinker [17] | 0.913±0.003 | 0.869±0.033 | 0.809±0.004 | 0.803±0.010 |
| | K-Paths [18] | 0.636±0.032 | 0.577±0.014 | 0.685±0.014 | 0.647±0.021 |
| | CAREPath | **0.981±0.003** | **0.943±0.011** | **0.929±0.010** | **0.915±0.013** |

Table S10: Performance comparison on Hetionet (Disease Cold-start Split).

| Methods | Models | AUROC | AUPRC | Accuracy | F1 |
|---|---|---|---|---|---|
| Graph-based | GraphSAGE [1] | 0.765±0.055 | 0.726±0.060 | 0.721±0.041 | 0.708±0.042 |
| | CompGCN [2] | 0.808±0.050 | 0.802±0.055 | 0.700±0.054 | 0.666±0.125 |
| | GIN [3] | 0.500±0.002 | 0.515±0.046 | 0.502±0.068 | 0.666±0.030 |
| | GAT [4] | 0.784±0.025 | 0.751±0.028 | 0.748±0.029 | 0.744±0.025 |
| KGE-based | TransE [5] | 0.691±0.038 | 0.678±0.027 | 0.504±0.041 | 0.666±0.038 |
| | TransR [6] | 0.530±0.059 | 0.561±0.066 | 0.512±0.046 | 0.646±0.047 |
| | RotatE [7] | 0.679±0.030 | 0.699±0.029 | 0.630±0.038 | 0.564±0.096 |
| | ComplEx [8] | 0.641±0.056 | 0.701±0.064 | 0.569±0.048 | 0.544±0.047 |
| | RESCAL [9] | 0.456±0.089 | 0.501±0.036 | 0.495±0.038 | 0.628±0.050 |
| LLM-based | BioBERT [10] | 0.841±0.011 | 0.795±0.007 | 0.766±0.011 | 0.752±0.012 |
| | BioLinkBERT [12] | 0.928±0.034 | 0.924±0.037 | 0.852±0.038 | 0.840±0.041 |
| | PubMedBERT [11] | 0.904±0.041 | 0.907±0.045 | 0.831±0.048 | 0.807±0.064 |
| | SapBERT [13] | 0.869±0.050 | 0.881±0.038 | 0.794±0.067 | 0.772±0.067 |
| Path-based | Node2Vec [14] | 0.850±0.051 | 0.837±0.005 | 0.748±0.046 | 0.696±0.065 |
| | DrugRep-KG [15] | 0.857±0.013 | 0.846±0.010 | 0.775±0.013 | 0.752±0.034 |
| | DREAMwalk [16] | 0.912±0.015 | 0.901±0.026 | 0.831±0.025 | 0.825±0.033 |
| | FuseLinker [17] | 0.922±0.003 | 0.927±0.013 | 0.872±0.013 | 0.860±0.010 |
| | K-Paths [18] | 0.913±0.005 | 0.907±0.021 | **0.907±0.010** | 0.891±0.003 |
| | CAREPath | **0.967±0.027** | **0.966±0.010** | 0.900±0.0.039 | **0.899±0.039** |

Table S11: Performance comparison on SuppKG (Disease Cold-start Split).

| Methods | Models | AUROC | AUPRC | Accuracy | F1 |
|---|---|---|---|---|---|
| Graph-based | GraphSAGE [1] | 0.820±0.017 | 0.895±0.003 | 0.774±0.018 | 0.834±0.015 |
| | CompGCN [2] | 0.942±0.004 | 0.873±0.002 | 0.879±0.003 | 0.914±0.002 |
| | GIN [3] | 0.933±0.005 | 0.923±0.088 | 0.863±0.011 | 0.897±0.009 |
| | GAT [4] | 0.702±0.022 | 0.760±0.022 | 0.778±0.008 | 0.850±0.007 |
| KGE-based | TransE [5] | 0.864±0.007 | 0.861±0.021 | 0.503±0.039 | 0.573±0.002 |
| | TransR [6] | 0.626±0.045 | 0.609±0.097 | 0.554±0.016 | 0.670±0.025 |
| | RotatE [7] | 0.742±0.017 | 0.772±0.025 | 0.535±0.045 | 0.670±0.038 |
| | ComplEx [8] | 0.834±0.019 | 0.849±0.016 | 0.632±0.069 | 0.428±0.116 |
| | RESCAL [9] | 0.723±0.010 | 0.698±0.034 | 0.503±0.040 | 0.632±0.48 |
| LLM-based | BioBERT [10] | 0.855±0.007 | 0.860±0.005 | 0.771±0.005 | 0.749±0.010 |
| | BioLinkBERT [12] | 0.903±0.005 | 0.908±0.012 | 0.822±0.005 | 0.818±0.010 |
| | PubMedBERT [11] | 0.899±0.006 | 0.904±0.011 | 0.818±0.007 | 0.811±0.014 |
| | SapBERT [13] | 0.895±0.008 | 0.902±0.013 | 0.816±0.008 | 0.814±0.014 |
| Path-based | Node2Vec [14] | 0.896±0.021 | 0.827±0.010 | 0.825±0.028 | 0.870±0.026 |
| | DrugRep-KG [15] | 0.910±0.005 | 0.841±0.011 | 0.811±0.034 | 0.856±0.006 |
| | DREAMwalk [16] | <u>0.922±0.007</u> | <u>0.960±0.004</u> | <u>0.858±0.012</u> | <u>0.898±0.009</u> |
| | FuseLinker [17] | 0.908±0.012 | 0.846±0.031 | 0.827±0.005 | 0.887±0.020 |
| | K-Paths [18] | 0.770±0.014 | 0.763±0.023 | 0.772±0.016 | 0.834±0.010 |
| | CAREPath | **0.967±0.002** | **0.970±0.012** | **0.906±0.004** | **0.905±0.004** |

Table S12: Performance comparison on KEGG50k (Disease Cold-start Split).

| Methods | Models | AUROC | AUPRC | Accuracy | F1 |
|---|---|---|---|---|---|
| Graph-based | GraphSAGE [1] | 0.843±0.030 | 0.694±0.061 | 0.795±0.027 | 0.620±0.072 |
| | CompGCN [2] | 0.878±0.045 | 0.743±0.066 | 0.656±0.005 | 0.644±0.007 |
| | GIN [3] | 0.823±0.052 | 0.745±0.050 | 0.810±0.022 | 0.617±0.063 |
| | GAT [4] | 0.863±0.022 | 0.743±0.050 | 0.804±0.022 | 0.633±0.062 |
| KGE-based | TransE [5] | 0.615±0.023 | 0.571±0.035 | 0.571±0.025 | 0.618±0.025 |
| | TransR [6] | 0.603±0.057 | 0.620±0.088 | 0.564±0.036 | 0.569±0.073 |
| | RotatE [7] | 0.633±0.014 | 0.660±0.043 | 0.520±0.045 | 0.647±0.033 |
| | ComplEx [8] | 0.639±0.020 | 0.682±0.027 | 0.544±0.038 | 0.671±0.038 |
| | RESCAL [9] | 0.557±0.020 | 0.561±0.043 | 0.508±0.042 | 0.502±0.046 |
| LLM-based | BioBERT [10] | 0.958±0.006 | 0.953±0.015 | 0.883±0.014 | 0.877±0.015 |
| | BioLinkBERT [12] | 0.976±0.002 | 0.975±0.003 | 0.920±0.003 | 0.918±0.006 |
| | PubMedBERT [11] | <u>0.977±0.005</u> | <u>0.976±0.003</u> | <u>0.922±0.006</u> | <u>0.920±0.007</u> |
| | SapBERT [13] | 0.950±0.012 | 0.951±0.006 | 0.873±0.023 | 0.867±0.019 |
| Path-based | Node2Vec [14] | 0.898±0.038 | 0.953±0.019 | 0.800±0.032 | 0.525±0.114 |
| | DrugRep-KG [15] | 0.952±0.003 | 0.977±0.003 | 0.891±0.012 | 0.881±0.007 |
| | DREAMwalk [16] | 0.921±0.029 | 0.870±0.035 | 0.819±0.029 | 0.598±0.081 |
| | FuseLinker [17] | 0.924±0.004 | 0.926±0.003 | 0.880±0.021 | 0.873±0.018 |
| | K-Paths [18] | 0.505±0.025 | 0.503±0.023 | 0.505±0.030 | 0.501±0.014 |
| | CAREPath | **0.989±0.002** | **0.985±0.002** | **0.950±0.005** | **0.950±0.005** |

**Drug Split(Drug Cold-start)**

Table S13: Performance comparison on MSI (Drug Cold-start Split).

| Methods | Models | AUROC | AUPRC | Accuracy | F1 |
|---|---|---|---|---|---|
| Graph-based | GraphSAGE [1] | 0.737±0.008 | 0.725±0.014 | 0.694±0.010 | 0.689±0.017 |
| | CompGCN [2] | 0.642±0.008 | 0.633±0.020 | 0.600±0.006 | 0.549±0.029 |
| | GIN [3] | 0.703±0.040 | 0.703±0.047 | 0.651±0.033 | 0.613±0.034 |
| | GAT [4] | 0.752±0.009 | 0.732±0.010 | 0.701±0.009 | 0.707±0.015 |
| KGE-based | TransE [5] | 0.648±0.020 | 0.657±0.010 | 0.593±0.021 | 0.442±0.039 |
| | TransR [6] | 0.576±0.015 | 0.571±0.024 | 0.551±0.009 | 0.449±0.058 |
| | RotatE [7] | 0.530±0.014 | 0.526±0.027 | 0.524±0.010 | 0.528±0.056 |
| | ComplEx [8] | 0.663±0.024 | 0.668±0.010 | 0.517±0.035 | 0.509±0.052 |
| | RESCAL [9] | 0.554±0.026 | 0.549±0.025 | 0.529±0.027 | 0.504±0.082 |
| LLM-based | BioBERT [10] | 0.806±0.021 | 0.804±0.018 | 0.743±0.021 | 0.736±0.024 |
| | BioLinkBERT [12] | 0.862±0.013 | 0.866±0.008 | 0.788±0.011 | 0.780±0.012 |
| | PubMedBERT [11] | 0.881±0.011 | 0.886±0.010 | 0.809±0.010 | 0.805±0.010 |
| | SapBERT [13] | 0.831±0.019 | 0.835±0.011 | 0.760±0.019 | 0.746±0.021 |
| Path-based | Node2Vec [14] | 0.864±0.012 | 0.877±0.010 | 0.799±0.009 | 0.786±0.009 |
| | DrugRep-KG [15] | 0.866±0.014 | 0.879±0.011 | 0.795±0.014 | 0.783±0.014 |
| | DREAMwalk [16] | 0.881±0.018 | 0.892±0.016 | 0.810±0.015 | 0.801±0.017 |
| | FuseLinker [17] | 0.896±0.010 | 0.917±0.024 | 0.832±0.020 | 0.814±0.009 |
| | K-Paths [18] | 0.891±0.009 | 0.902±0.013 | 0.835±0.012 | 0.825±0.010 |
| | CAREPath | **0.918±0.005** | **0.924±0.005** | **0.843±0.005** | **0.838±0.008** |

Table S14: Performance comparison on PrimeKG (Drug Cold-start Split).

| Methods | Models | AUROC | AUPRC | Accuracy | F1 |
|---|---|---|---|---|---|
| Graph-based | GraphSAGE [1] | 0.942±0.011 | 0.910±0.009 | 0.889±0.008 | 0.872±0.010 |
| | CompGCN [2] | 0.944±0.009 | 0.925±0.007 | 0.877±0.008 | 0.860±0.009 |
| | GIN [3] | 0.746±0.119 | 0.668±0.136 | 0.707±0.107 | 0.519±0.305 |
| | GAT [4] | 0.956±0.008 | 0.938±0.007 | 0.904±0.006 | 0.889±0.008 |
| KGE-based | TransE [5] | 0.792±0.033 | 0.748±0.055 | 0.620±0.020 | 0.603±0.042 |
| | TransR [6] | 0.511±0.082 | 0.497±0.091 | 0.480±0.074 | 0.464±0.095 |
| | RotatE [7] | 0.654±0.011 | 0.501±0.035 | 0.573±0.015 | 0.550±0.047 |
| | ComplEx [8] | 0.857±0.066 | 0.857±0.049 | 0.604±0.024 | 0.679±0.049 |
| | RESCAL [9] | 0.733±0.041 | 0.654±0.025 | 0.575±0.026 | 0.572±0.003 |
| LLM-based | BioBERT [10] | 0.966±0.004 | 0.954±0.008 | 0.902±0.006 | 0.884±0.012 |
| | BioLinkBERT [12] | 0.948±0.007 | 0.928±0.015 | 0.868±0.009 | 0.843±0.013 |
| | PubMedBERT [11] | 0.958±0.005 | 0.944±0.011 | 0.883±0.010 | 0.861±0.015 |
| | SapBERT [13] | 0.918±0.011 | 0.891±0.021 | 0.831±0.013 | 0.789±0.020 |
| Path-based | Node2Vec [14] | **0.969±0.006** | **0.964±0.005** | **0.913±0.004** | **0.891±0.006** |
| | DrugRep-KG [15] | 0.704±0.030 | 0.749±0.053 | 0.664±0.035 | 0.718±0.047 |
| | DREAMwalk [16] | **0.969±0.005** | **0.964±0.005** | 0.911±0.008 | 0.889±0.011 |
| | FuseLinker [17] | 0.909±0.004 | 0.865±0.024 | 0.851±0.014 | 0.840±0.005 |
| | K-Paths [18] | 0.601±0.018 | 0.579±0.032 | 0.554±0.015 | 0.550±0.010 |
| | CAREPath | 0.957±0.005 | 0.943±0.007 | 0.861±0.014 | 0.817±0.021 |

Table S15: Performance comparison on Hetionet (Drug Cold-start Split).

| Methods | Models | AUROC | AUPRC | Accuracy | F1 |
|---|---|---|---|---|---|
| Graph-based | GraphSAGE [1] | 0.852±0.013 | 0.816±0.024 | 0.767±0.032 | 0.760±0.043 |
| | CompGCN [2] | 0.809±0.012 | 0.803±0.018 | 0.720±0.016 | 0.729±0.034 |
| | GIN [3] | 0.562±0.123 | 0.559±0.119 | 0.554±0.098 | 0.553±0.279 |
| | GAT [4] | 0.884±0.026 | 0.856±0.037 | 0.819±0.022 | 0.822±0.021 |
| KGE-based | TransE [5] | 0.523±0.056 | 0.520±0.117 | 0.482±0.094 | 0.642±0.095 |
| | TransR [6] | 0.656±0.021 | 0.650±0.025 | 0.575±0.036 | 0.671±0.024 |
| | RotatE [7] | 0.539±0.030 | 0.532±0.042 | 0.537±0.033 | 0.511±0.055 |
| | ComplEx [8] | 0.723±0.032 | 0.723±0.023 | 0.646±0.025 | 0.598±0.040 |
| | RESCAL [9] | 0.647±0.034 | 0.640±0.018 | 0.589±0.036 | 0.588±0.043 |
| LLM-based | BioBERT [10] | 0.871±0.008 | 0.877±0.015 | 0.793±0.010 | 0.788±0.016 |
| | BioLinkBERT [12] | 0.936±0.016 | 0.933±0.016 | 0.867±0.025 | 0.864±0.021 |
| | PubMedBERT [11] | 0.936±0.021 | 0.936±0.022 | 0.867±0.026 | 0.865±0.029 |
| | SapBERT [13] | 0.914±0.014 | 0.913±0.021 | 0.835±0.021 | 0.823±0.036 |
| Path-based | Node2Vec [14] | 0.916±0.009 | 0.911±0.012 | 0.834±0.014 | 0.830±0.020 |
| | DrugRep-KG [15] | <u>0.957±0.005</u> | **0.979±0.004** | **0.899±0.008** | **0.928±0.008** |
| | DREAMwalk [16] | 0.842±0.044 | 0.844±0.047 | 0.723±0.044 | 0.655±0.069 |
| | FuseLinker [17] | 0.973±0.006 | <u>0.972±0.004</u> | 0.893±0.017 | <u>0.880±0.012</u> |
| | K-Paths [18] | 0.955±0.003 | 0.944±0.007 | 0.872±0.012 | 0.864±0.010 |
| | CAREPath | **0.962±0.008** | 0.961±0.010 | <u>0.897±0.011</u> | 0.897±0.014 |

Table S16: Performance comparison on SuppKG (Drug Cold-start Split).

| Methods | Models | AUROC | AUPRC | Accuracy | F1 |
|---|---|---|---|---|---|
| Graph-based | GraphSAGE [1] | 0.842±0.012 | 0.904±0.010 | 0.787±0.020 | 0.841±0.019 |
| | CompGCN [2] | 0.943±0.004 | 0.974±0.002 | 0.881±0.005 | 0.915±0.004 |
| | GIN [3] | 0.931±0.009 | 0.965±0.005 | 0.865±0.012 | 0.899±0.009 |
| | GAT [4] | 0.716±0.027 | 0.775±0.031 | 0.773±0.019 | 0.846±0.014 |
| KGE-based | TransE [5] | 0.866±0.011 | 0.871±0.010 | 0.656±0.019 | 0.578±0.004 |
| | TransR [6] | 0.725±0.027 | 0.725±0.024 | 0.512±0.019 | 0.670±0.015 |
| | RotatE [7] | 0.491±0.010 | 0.481±0.024 | 0.504±0.016 | 0.631±0.019 |
| | ComplEx [8] | 0.862±0.009 | 0.872±0.006 | 0.787±0.014 | 0.764±0.018 |
| | RESCAL [9] | 0.758±0.018 | 0.725±0.031 | 0.501±0.019 | 0.638±0.006 |
| LLM-based | BioBERT [10] | 0.899±0.005 | 0.904±0.011 | 0.821±0.005 | 0.816±0.011 |
| | BioLinkBERT [12] | 0.891±0.011 | 0.897±0.016 | 0.812±0.012 | 0.805±0.017 |
| | PubMedBERT [11] | 0.898±0.011 | 0.905±0.013 | 0.818±0.015 | 0.811±0.019 |
| | SapBERT [13] | 0.867±0.011 | 0.874±0.017 | 0.786±0.011 | 0.776±0.018 |
| Path-based | Node2Vec [14] | 0.933±0.004 | <u>0.968±0.003</u> | <u>0.870±0.006</u> | **0.908±0.005** |
| | DrugRep-KG [15] | 0.880±0.013 | 0.912±0.013 | 0.821±0.024 | 0.843±0.008 |
| | DREAMwalk [16] | <u>0.917±0.016</u> | 0.959±0.007 | 0.853±0.019 | 0.894±0.014 |
| | FuseLinker [17] | 0.881±0.005 | 0.906±0.018 | 0.833±0.013 | 0.852±0.017 |
| | K-Paths [18] | 0.793±0.012 | 0.800±0.032 | 0.732±0.025 | 0.755±0.029 |
| | CAREPath | **0.967±0.002** | **0.970±0.002** | **0.907±0.002** | <u>0.905±0.002</u> |

Table S17: Performance comparison on KEGG50k (Drug Cold-start Split).

| Methods | Models | AUROC | AUPRC | Accuracy | F1 |
|---|---|---|---|---|---|
| Graph-based | GraphSAGE [1] | 0.942±0.003 | 0.858±0.010 | 0.889±0.008 | 0.825±0.0115 |
| | CompGCN [2] | 0.879±0.005 | 0.757±0.013 | 0.656±0.013 | 0.644±0.014 |
| | GIN [3] | 0.959±0.006 | 0.909±0.010 | 0.929±0.004 | 0.887±0.006 |
| | GAT [4] | 0.960±0.006 | 0.908±0.009 | 0.926±0.002 | 0.883±0.005 |
| KGE-based | TransE [5] | 0.750±0.026 | 0.748±0.035 | 0.657±0.028 | 0.704±0.022 |
| | TransR [6] | 0.691±0.020 | 0.701±0.015 | 0.601±0.048 | 0.653±0.026 |
| | RotatE [7] | 0.571±0.011 | 0.543±0.016 | 0.522±0.012 | 0.663±0.006 |
| | ComplEx [8] | 0.834±0.053 | 0.855±0.044 | 0.693±0.046 | 0.558±0.089 |
| | RESCAL [9] | 0.687±0.061 | 0.665±0.047 | 0.507±0.010 | 0.578±0.018 |
| LLM-based | BioBERT [10] | 0.972±0.003 | 0.968±0.006 | 0.918±0.008 | 0.917±0.011 |
| | BioLinkBERT [12] | 0.977±0.004 | 0.975±0.006 | 0.924±0.009 | 0.923±0.013 |
| | PubMedBERT [11] | 0.979±0.005 | 0.978±0.007 | 0.929±0.011 | 0.928±0.014 |
| | SapBERT [13] | 0.958±0.002 | 0.957±0.006 | 0.890±0.007 | 0.887±0.013 |
| Path-based | Node2Vec [14] | 0.977±0.006 | 0.963±0.005 | <u>0.951±0.002</u> | 0.919±0.004 |
| | DrugRep-KG [15] | <u>0.985±0.002</u> | <u>0.982±0.004</u> | <u>0.951±0.012</u> | <u>0.931±0.003</u> |
| | DREAMwalk [16] | 0.984±0.007 | 0.973±0.010 | **0.957±0.011** | 0.929±0.017 |
| | FuseLinker [17] | 0.953±0.005 | 0.941±0.004 | 0.910±0.002 | 0.908±0.003 |
| | K-Paths [18] | 0.583±0.050 | 0.562±0.046 | 0.520±0.017 | 0.501±0.078 |
| | CAREPath | **0.987±0.001** | **0.985±0.001** | 0.942±0.005 | **0.941±0.005** |

#### S6. Supplementary Case Studies

**Dermatitis, Atopic → IL6 → NR3C1 → Prednisolone**

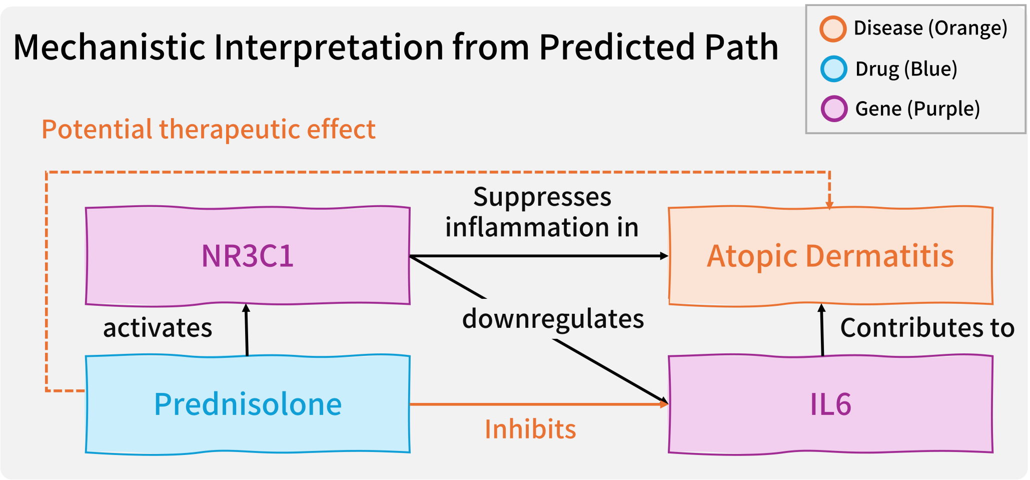


Fig. S1: **Mechanistic interpretation for a clinically established therapy.** CAREPath retrieves a mechanistic chain for atopic dermatitis, highlighting that prednisolone activates NR3C1 (glucocorticoid receptor), which suppresses inflammation and downregulates IL6, a cytokine contributing to inflammatory pathology. The dashed outline denotes the predicted potential therapeutic effect supported by the inferred path.

We provide an additional qualitative case study illustrating that CAREPath can recover clinically used therapies with an explicit mechanistic rationale. As shown in Fig.S1, CAREPath links atopic dermatitis to prednisolone through the IL6–NR3C1 axis: IL6 contributes to inflammatory pathology in atopic dermatitis, while prednisolone activates NR3C1 to suppress inflammatory programs and downregulate IL6.

### References

[1] Will Hamilton, Zhitao Ying, and Jure Leskovec. Inductive representation learning on large graphs. *Advances in neural information processing systems*, 30, 2017.

[2] Shikhar Vashishth, Soumya Sanyal, Vikram Nitin, and Partha Talukdar. Composition-based multi-relational graph convolutional networks. *arXiv preprint arXiv:1911.03082*, 2019.

[3] Keyulu Xu, Weihua Hu, Jure Leskovec, and Stefanie Jegelka. How powerful are graph neural networks? *arXiv preprint arXiv:1810.00826*, 2018.

[4] Petar Veličković, Guillem Cucurull, Arantxa Casanova, Adriana Romero, Pietro Lio, and Yoshua Bengio. Graph attention networks. *arXiv preprint arXiv:1710.10903*, 2017.

[5] Antoine Bordes, Nicolas Usunier, Alberto Garcia-Duran, Jason Weston, and Oksana Yakhnenko. Translating embeddings for modeling multi-relational data. *Advances in neural information processing systems*, 26, 2013.

[6] Yankai Lin, Zhiyuan Liu, Maosong Sun, Yang Liu, and Xuan Zhu. Learning entity and relation embeddings for knowledge graph completion. In *Proceedings of the AAAI conference on artificial intelligence*, volume 29, 2015.

[7] Zhiqing Sun, Zhi-Hong Deng, Jian-Yun Nie, and Jian Tang. Rotate: Knowledge graph embedding by relational rotation in complex space. *arXiv preprint arXiv:1902.10197*, 2019.

[8] Théo Trouillon, Johannes Welbl, Sebastian Riedel, Éric Gaussier, and Guillaume Bouchard. Complex embeddings for simple link prediction. In *International conference on machine learning*, pages 2071–2080. PMLR, 2016.

[9] Maximilian Nickel, Volker Tresp, Hans-Peter Kriegel, et al. A three-way model for collective learning on multi-relational data. In *Icml*, volume 11, pages 3104482–3104584, 2011.

[10] Jinhyuk Lee, Wonjin Yoon, Sungdong Kim, Donghyeon Kim, Sunkyu Kim, Chan Ho So, and Jaewoo Kang. Biobert: a pre-trained biomedical language representation model for biomedical text mining. *Bioinformatics*, 36(4):1234–1240, 2020.

[11] Yu Gu, Robert Tinn, Hao Cheng, Michael Lucas, Naoto Usuyama, Xiaodong Liu, Tristan Naumann, Jianfeng Gao, and Hoifung Poon. Domain-specific language model pretraining for biomedical natural language processing. *ACM Transactions on Computing for Healthcare (HEALTH)*, 3(1):1–23, 2021.

[12] Michihiro Yasunaga, Jure Leskovec, and Percy Liang. Linkbert: Pretraining language models with document links. *arXiv preprint arXiv:2203.15827*, 2022.

[22] Minoru Kanehisa, Miho Furumichi, Mao Tanabe, Yoko Sato, and Kanae Morishima. Kegg: new perspectives on genomes, pathways, diseases and drugs. *Nucleic acids research*, 45(D1):D353–D361, 2017.

[23] Zahra Alaeddini, Parham Moradi, and Bahram Sadeghi Bigham. A graph-based knowledge distillation framework for drug repurposing via multi-task learning. *Engineering Applications of Artificial Intelligence*, 162:112752, 2025.
